# A human stem cell-derived neurosensory-epithelial circuitry on-a-chip to model herpes simplex virus reactivation

**DOI:** 10.1101/2020.11.19.390260

**Authors:** Pietro Giuseppe Mazzara, Elena Criscuolo, Marco Rasponi, Luca Massimino, Sharon Muggeo, Cecilia Palma, Matteo Castelli, Massimo Clementi M, Roberto Burioni, Nicasio Mancini, Vania Broccoli, Nicola Clementi

## Abstract

Both emerging viruses and well-known viral pathogens endowed with neurotropism can either impair directly the neuronal functions or induce physio-pathological changes by diffusing from the periphery through neurosensory-epithelial connections. However, the current lack of an *in vitro* system modeling the connectivity between human neurons and peripheral tissues excludes the analysis of viral latency and reactivation and the assessment of natural/artificial induced anti-viral immunity. In this study, we developed the first stable topographic neurosensory-epithelial connection on-a-chip using human stem cell derived dorsal root ganglia (DRG) sensory neurons. Bulk and single cell transcriptomics showed that different combinations of key receptors for Herpes Simplex Virus 1 (HSV-1) are expressed by each sensory neuronal cell type. This neuronal-epithelial circuitry enabled a detailed analysis of the HSV infectivity faithfully modeling its dynamics and cell type specificity. The reconstitution of an organized connectivity between human sensory neurons and keratinocytes into microfluidic chips provides for the first time a powerful *in vitro* platform to model viral latency and reactivation of human viral pathogens.

## Introduction

Herpes Simplex Viruses (HSV) have the characteristic of establishing long-term latency into peripheral neurons, from which reactivate causing symptoms in innervated tissues. The frequency and severity of reactivations vary according to the characteristics of the host’s immune response and of the virus isolate involved (White et al. 2012), but the essential molecular mechanisms have not yet been clarified (Nicoll et al. 2012). Indeed, reactivations studies of neurotropic viruses are a concern. *In vivo* experiments have been described, but they are endowed with the limitations of using a human-adapted pathogen in animal model systems, often resulting in completely different pathophysiology (Sawtell & Thompson 1992; Birmanns et al. 1993; Chiuppesi et al. 2012; Yao et al. 2014; BenMohamed et al. 2016; Ramakrishna et al. 2015; Berges & Tanner 2014). Moreover, these studies required high numbers while leading to results not entirely reflecting the human condition. Similarly, no satisfactory result has been achieved *in vitro*.

The first aspect to be addressed is a convenient neuronal system for a reliable HSV reactivation protocol. Different approaches have been described so far, but no procedure was accepted unanimously. HSV infection has been tested in several experimental settings (Sun et al. 2016; Pourchet et al. 2017; Y. Liu et al. 2014; Thellman & Triezenberg 2017; Danaher, Jacob & Miller 1999), using differentiated and undifferentiated human neuroblastoma-derived cell lines, and subsequent reactivation using thermal or chemical stimuli. Furthermore, complex systems have been developed using human induced pluripotent stem cell (hiPSC)-derived neurons (Lee et al. 2012). It has been observed that, while useful to study the dynamics of viral expression, these cultures do not reproduce all the complexities of HSV-1 reactivation *in vivo*. Following the advent of 3D cellular cultures, designated organoids, these models have also been investigated (D’Aiuto et al. 2019). In particular, for brain organoids, the 3D culture systems mimic aspects of brain architecture and allow complex cell-cell communication, cell-cell interaction, and cell-extracellular matrix interaction. Brain organoids have given a great impulse to the study of other neurotropic viruses, including Zika virus and VZV (Xu et al. 2019; Sloutskin et al. 2013). Recent data describes how they can be efficiently infected by HSV-1, but the treatment of latently infected organoids with several stimuli has not led to viral reactivation (D’Aiuto et al. 2019).

The second issue is to reconstitute the neural connection with the cells affected by productive infection to allow axonal transport and neuron-to-cell spread. This issue is crucial for studying HSV infection. Only an *in vitro* method has been developed, using pseudorabies virus (PRV) (W. W. Liu et al. 2008). The method centers on a microfluidic chamber system that directs the growth of axons into a fluidically isolated environment and recapitulates all known aspects of neuron-to-cell spread. Microfluidic chip devices indeed have been rapidly improved. Their high versatility and customization allows the reproduction of specific environments that mimic the differences between human tissue compartments, to dissect physiological conditions that could only be analyzed *in vivo* (Iannielli et al. 2019). Results were achieved on VZV reactivation from latency by using hESC-derived neurons, but no *neuron-to-cell* connection was investigated (Markus et al. 2015). Therefore, induction and reactivation from latency of herpesviruses on human DRGO through infection of primary keratinocytes sinaptically connected with them was not described yet. In this study, we took advantage of human iPSC-derived 3D dorsal root ganglion-like organoids (DRGOs), generated by *in vitro* differentiation of hiPSCs in 3D cultures (Mazzara et al. 2020). Molecular and genomic analyses and topographic connectivity to peripheral targets have confirmed that DRGOs are populated by nociceptive, mechanoreceptive and proprioceptive neurons. Herein, we have moved forward by establishing a fully human circuit between DRGO-derived sensory neurons and primary keratinocytes. Furthermore, the circuit was organized in a patterned topography within a miniaturized microfluidics platform, thus making it extremely feasible for its reproducibility and translational application.

## Results

### Setting the neuro-epithelial co-culture model in a patterned microfluidic chip culture system

A thorough analysis of virus reactivation requires a humanized system where keratinocytes and neurons can be simultaneously cultured for extensive time developing direct connectivity. To this aim we decided to evaluate the possibility to develop a new neuron-keratinocyte co-culture system using hiPSC-derived dorsal root ganglia organoids (DRGOs) that we have recently derived and showed to generate molecularly-defined peripheral sensory neurons capable of making specific contacts with their respective target tissues (Mazzara et al. 2020). DRGOs were obtained by differentiating transgene-free hiPSC-derived aggregates in 3D culture conditions and sequentially exposed to small molecules to induce peripheral nervous system identity (**Figure S1A**). This protocol led to the generation of rounded spheroids emitting radially projecting axonal projections (**Figure S1B**) which can be maintained in vitro for several weeks. Successful commitment to peripheral sensory neuronal identity was shown by high expression levels of specific molecular markers including *BRN3a* and *NAV1.7* (**Figure S1C**). Triple immunostaining identified peripheral sensory neurons expressing different combinations of TRK receptors confirming their correct differentiation in nociceptive (TRKA+/TRKC−), mechanoreceptive (TRKB+/TRKC+) and proprioceptive (TRKB−/TRKC+) neurons (**Figure S1D-F**). Bulk and single-cell RNA-seq confirmed and extended this analysis showing that DRGOs include also Satellite cells and Schwann cell progenitors, mirroring the overall cell composition of somatic DRGs (Mazzara et al. 2020).

For establishing co-cultures, either immortalized (HaCaT) or primary (NHEK) keratinocytes were seeded on coated slides, and 24h later DRGOs were added on the monolayers. Different culture media were tested to find the correct composition to meet the nutritional requirements of all the cell types. Unfortunately, all tested conditions were unable to ensure proper cell viability as shown by their morphology (**Figure S2A**). DRGO *in vitro* culture was incompatible with HaCaT medium, even when diluted 1:2 most of DRGOs were unable to attach to the plate and contact their target epithelial cells. The remaining DRGOs, capable to adhere to the substrate, appeared completely degenerated only after few days. In NHEK medium, DRGO neurons showed a better survival, but their differentiation was compromised since they failed to develop and extend axonal projections (Chalazonitis et al. 2001). On the other hand, both keratinocyte cultures showed an intact monolayer with proper cell morphology when cultured with their medium supplemented with BDNF, GDNF, NGF, NT-3 and AA. However, both NHEK and HaCaT cells cannot survive in DRGO medium promptly losing their monolayer cell organization before a generalized loss in cell survival. This drastic susceptibility to medium composition was confirmed by cell proliferation assay (**Figure S2B**). Thus, direct co-culture between neurons and keratinocytes remained unfeasible due to inherently different culture condition requirements.

In order to overcome the direct co-culture limitations, we sought to reconstitute the co-culture system into an appropriate microfluidic chip to prevent media mixing. We generated a custom-designed chip composed of two chambers, one for each cellular type, connected together by microgrooves to allow axonal projections to reach and connect with the keratinocytes (**Figure 1A**). Long term cell survival and proper cell morphology was detected in both chambers providing evidence that each culture medium was fluidically isolated in its chamber, with no evident morphological changes in the second cell type within the same chip (**Figure 1B**). However, when considering neuron-to-target connections, there were obvious differences between chips seeded with HaCaT or NHEK cells. Although all cultured cells were healthy, connections were observed only if primary keratinocytes were present. Axonal terminals were degenerate when they reached the HaCaT chamber, suggesting that the serum necessary for their growth negatively interfered with normal neuronal behavior, similarly to what we observed in previous direct co-cultures. Microfluidic devices are a convenient tool for direct imaging studies on isolated neurons, as well as seeded cells. Immunofluorescence analysis confirmed the innervation of the keratinocyte with multiple DRGO axons and the formation of stable connections between DRGOs and NHEKs. Indeed, we clearly observed Synapsin and vGlut positive DRGO free nerve endings on NHEKs closely recapitulating the typical anatomical configuration of the nociceptive contacts within the skin keratinocytes *in vivo* (**Figure 1C**). These results confirmed that the design of the microfluidic chip with an array of microgrooves connecting the two lateral chambers was efficient in preventing culture media mixing, while enabling axonal connectivity between the two compartments. Additionally, we provided evidence that DRGOs are a suitable model for multiorgan studies *in vitro*, reconstituting stable and topographic neuronal circuitries with different peripheral tissue targets.

**Figure 1.**
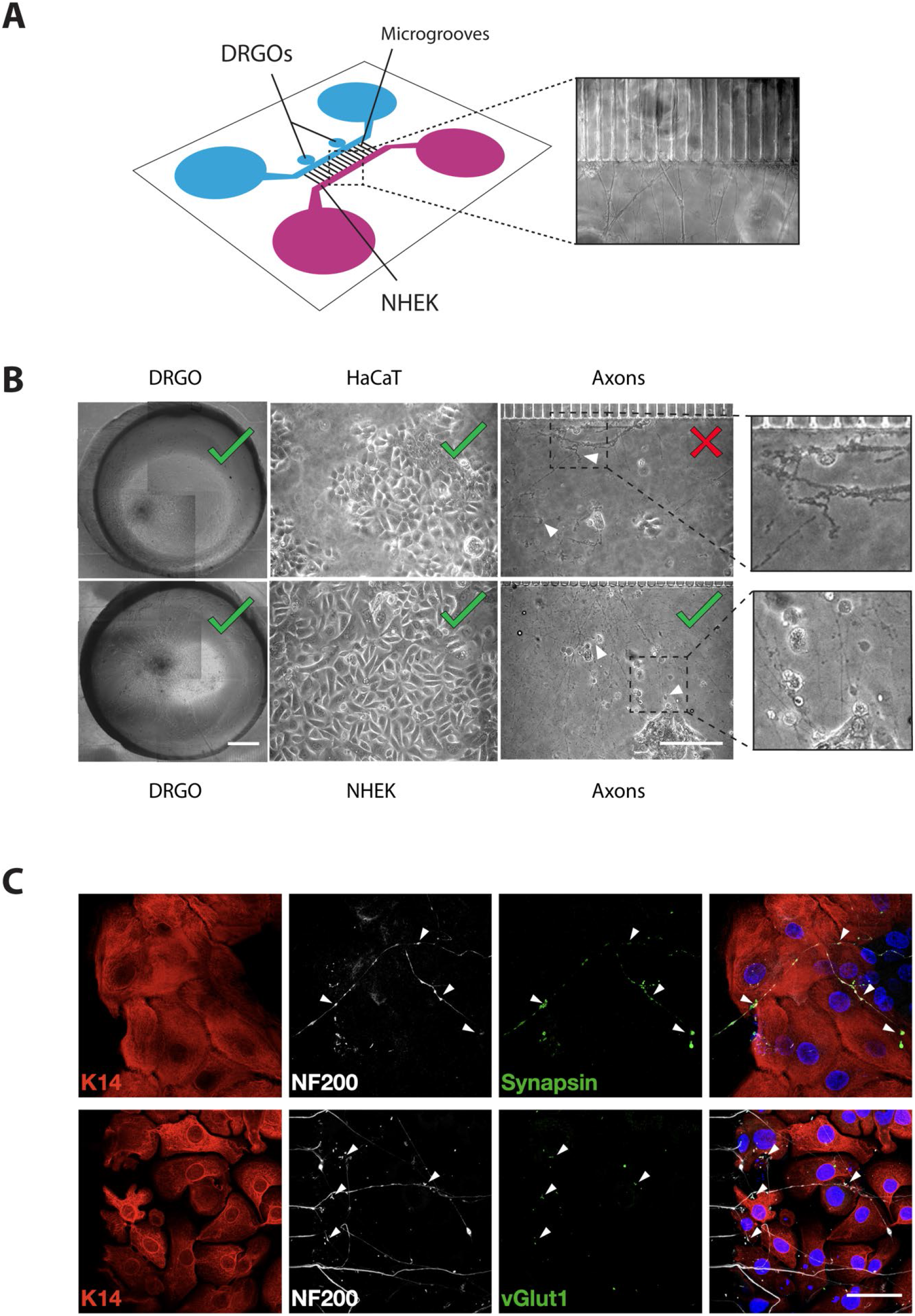
Co-culture of DRGOs and keratinocytes on-a-chip. **(A)** Graphical representation of microfluidics chip and bright-field image (20x magnification) of axonal extensions from DRGOs chamber to NHEK chamber through microgrooves. **(B)** Bright-field microscopy images of cells co-cultured in microfluidic chips, with neuron-to-cell connection (white triangles). Scale bars, DRGO 200 μm, HaCaT/NHEK/Axons 100 μm. **(C)** Immunofluorescence analysis of free nerve ending neuron-to-cell connection using K14 (red), NF200 (white), Synapsin (green, up) and vGlut1 (green, down). Total nuclear DNA is counterstained with Hoechst (blue). Scale bar, 50 μm.

### DRGO gene expression competence to HSV-1 infection

It is well known that HSV-1 entry mechanism is a highly complex process involving different molecular components of its cellular targets (Agedelis & Shukla, 2015). To confirm the DRGO sensory neurons competence for HSV infectivity, we next evaluated the main HSV cellular receptors expression in DRGOs. To this end, we carried out a computational analysis on global and single cell RNA-sequencing datasets on isolated DRGOs generated in our previous work (Mazzara et al. 2020). When compared to iPSCs at global transcriptomic level, we confirmed that DIV40 DRGOs and primary DRGs are closely related with respect to the expression of the main genes involved in virus-cell contact, in particular the main HSV-1 receptor *NECTIN1*, but also *TNFRSF14* (*HVEM*), PILRA, *NRP1*, *MYH9*, *ITGA5*, *ITGA8*, *ITGB6* and *ITGB8* (**Figure 2A**). Interestingly, this similarity is not strikingly related to HSV-1 receptors, but is a more general feature of DRGOs, underlined from the close relation to primary DRG in regard Heparan sulfate biosynthetic pathways and, more in general, integrins (**Figure 2B-C**). Remarkably, such similarities between primary and stem cell derived ganglion neurons were evident even if DIV40 DRGO neurons are generally not completely mature yet (Mazzara et al. 2020), and some differences are still present, such as for *MAG*, *HS3ST3B1* and *SCD1* (**Figure 2A)**.

**Figure 2.**
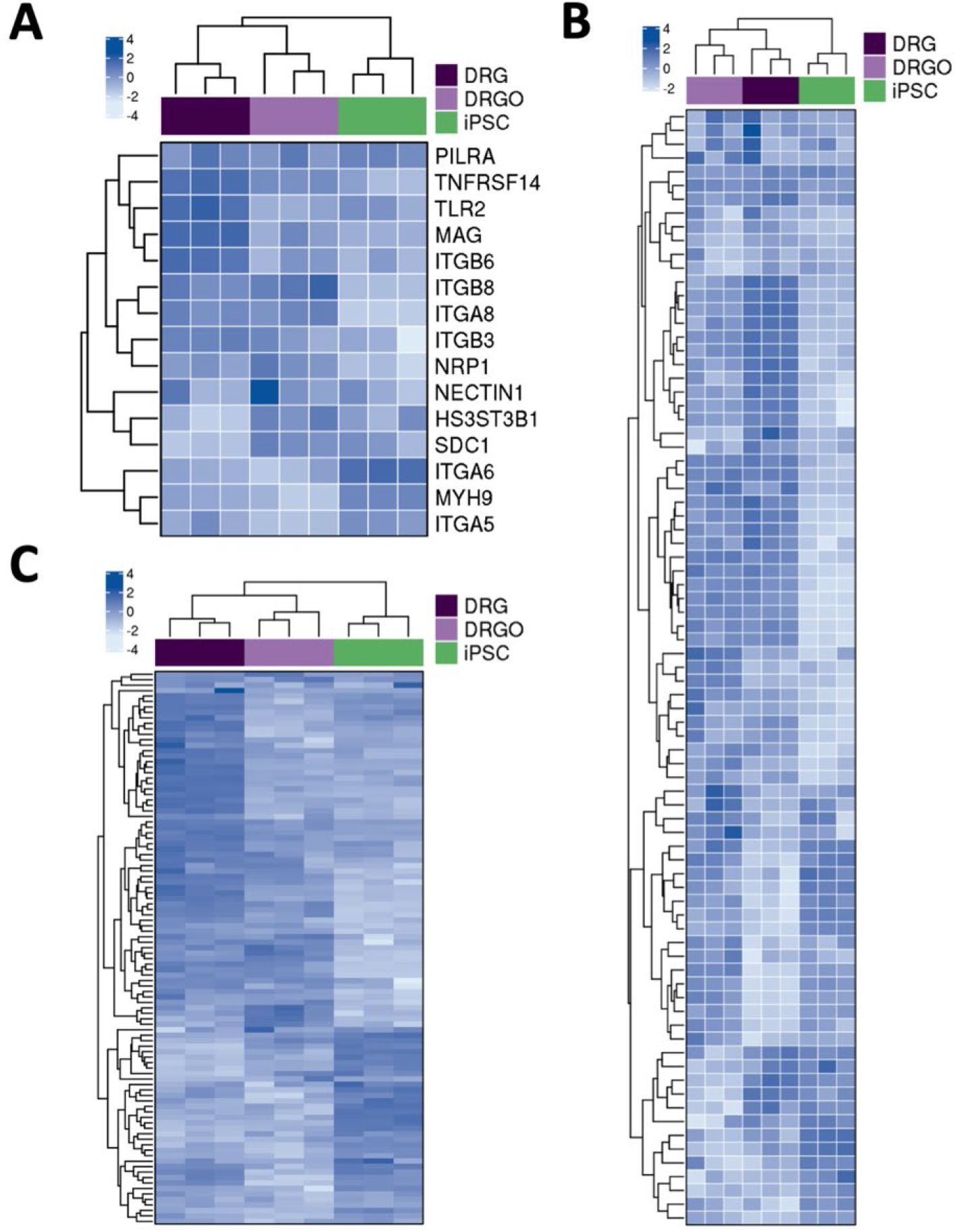
Global gene expression profile of HSV receptors in DRGO. (**A**) Supervised gene expression heatmaps showing the HSV receptors genes differentially expressed either in DRG vs iPSC, or DIV 40 DRGO vs iPSC; the correlation between samples is also shown as an unsupervised hierarchical clustered dendrogram on the side. (**B** and **C**) Gene expression heatmaps showing the differentially expressed genes belonging to aggregated GO categories associated with Heparan sulfate biosynthetic pathways (**B**) and integrin mediated signaling pathway (**C**); the correlation between samples is also shown as an unsupervised hierarchical clustered dendrogram on the side.

To better understand the cellular diversity of DRGO receptor composition in more mature DRGOs, we analyzed scRNA-seq datasets performed on 5,363 single cells isolated from two independent batches of DIV80 DRGOs, from which we identified 14 cell clusters based on their overall gene signature, containing all three sensory neuron subtypes (nociceptive, mechanoreceptive and proprioceptive neurons), satellite cells, Schwann cells (**Figure 3A**), fully recapitulating the cellular composition of somatic DRGs. Interestingly, we observed different combinations of expressed HSV receptors in the different DRGO sensory neuronal and glial cells (**Figure 3B-J**). All neuronal populations (C3, C4, C8) exhibited *NECTIN1* and *NRP1* expression, while in glial cells (C9-C13) these receptors were almost absent. The nociceptor cluster (C4) presented, differently from proprioceptors, high levels of *SDC1* and *HS3ST3B1*. The mechanoceptor cluster (C8) was the only neuronal group expressing *ITGB8* and, together with Schwann cells, *ITGA6*. In glial cells, the satellite cell clusters (C9-C11) were characterized by low levels of *HS3ST3B1* and *ITGA6*, and Schwann cell clusters (C12, C13) present high level of *ITGA6* and low level of *ITGB8* (**Figure 3B-J**). Collectively, these findings demonstrate that DRGOs resemble primary DRGs with respect to the global expression of HSV receptors and binding factors. However, we revealed a remarked cell-type specific expression pattern of important components of the viral-to-host interaction. These findings suggest that each single sensory neuronal population might have a specific profile of its interaction with HSV, and DRGOs might represent a convenient system which recapitulates the full complexity of the DRG cellular system.

**Figure 3.**
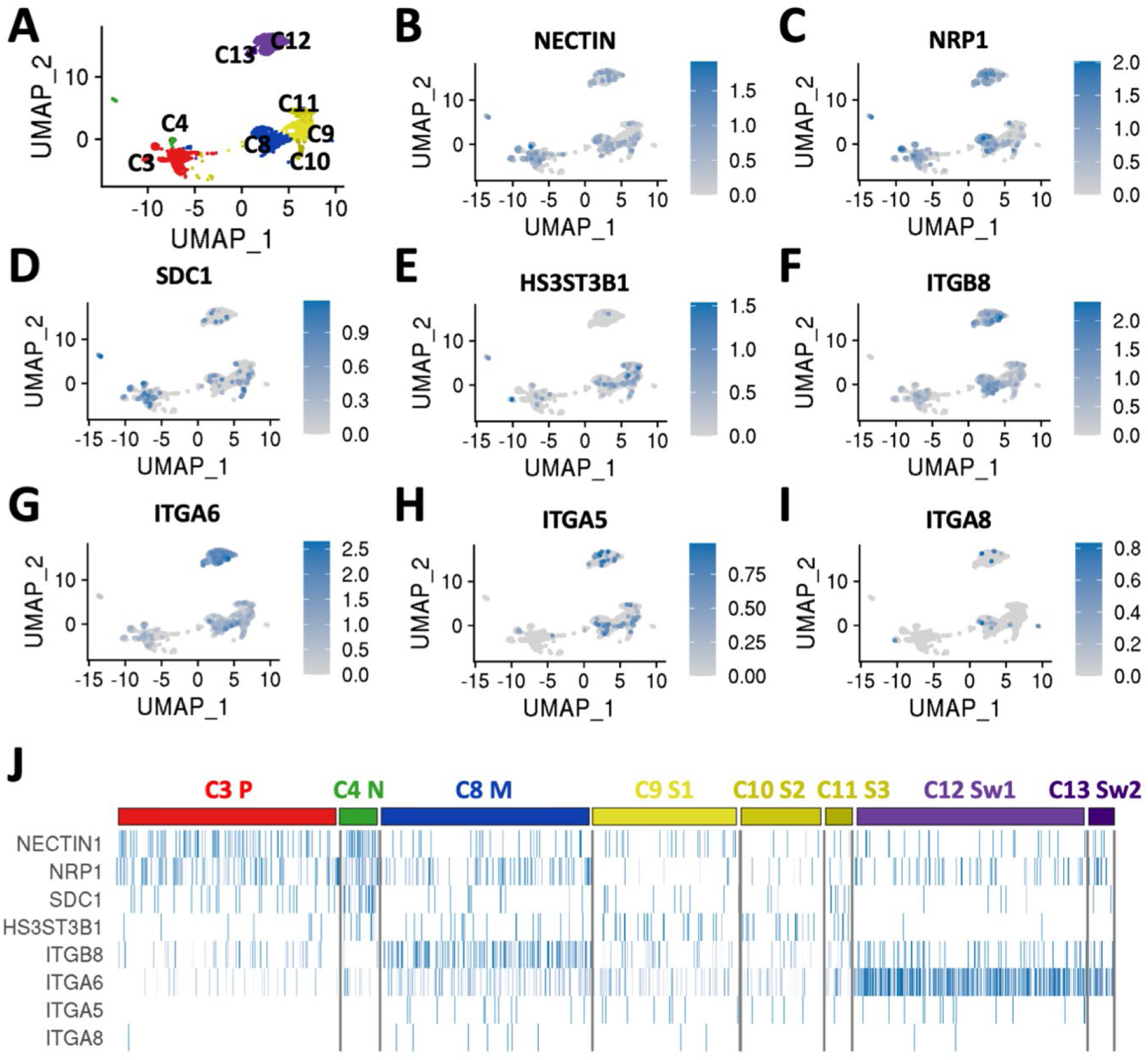
Single cell transcriptional profile of HSV receptors in DRGO. (**A**) Uniform Manifold Approximation and Projection (UMAP) plot displaying multidimensional reduction and clustering of single cell RNA-Seq data from DIV 80 DRGOs showing clusters of mature sensory neurons (proprioceptors C3, nociceptors C4, Mechanoceptors C8), satellite cells C9-C11, and Schwann cells C12, C13 (modified from Mazzara et al., in press). (**B-I**) UMAP plots highlighting normalized expression values of *NECTIN1* (**B**), *NRP1* (**C**), *SDC1* (**D**), *HS3ST3B1* (**E**), *ITGB8* (**F**), *ITGA6* (**G**), *ITGA5* (**H**), and *ITGA8* (**I**). (**J**) Heatmap showing normalized expression values of cell lineage-specific genes within the different clusters.

### Validation of the neuro-epithelial microfluidic platform for viral infectivity studies

We, next, asked to what extent this topographic human neuronal-epithelial circuitry can be informative for study the complex behavior of HSV infectivity. At first, we asked if virus particles were blocked in their passive diffusion between chambers, even during lytic infection when there is wide viral spreading among cells and massive production of new virions. This is of primary importance to validate our model for studies of neuron-to-cell HSV diffusion. To follow the lytic and latent phases of HSV infection, a recombinant HSV-1 KOS was used in the present study, incorporating enhanced green fluorescent protein (EGFP) driven by the viral promoters ICP0 and monomeric red fluorescent protein (RFP) driven by the viral promoters glycoprotein C (gC) as reporters for infected cells or cells in lytic phase, respectively (Decman et al. 2005). In detail, LAT is the only actively transcribed gene from the virus in latently infected cells, and the LAT region is located between two early lytic genes, ICP0 and ICP27. The mLAT transcript extends approximately 8.5 kb and ICP0 is transcribed from the opposite strand and overlaps with the mLAT transcribed region (Q. Chen et al. 2007). Based on the ChIP studies (Kubat, Tran, et al. 2004; Kubat, Amelio, et al. 2004), the boundary between transcription-permissive and -nonpermissive regions is located near the 3′ end of the ICP0, which overlaps with the LAT 2-kb intron by about 700 bp (Amelio et al. 2006). Thus, the ICP0 promoter is actively transcribed during latency, whereas the surrounding regions, such as the lytic-specific gene ICP0, are silenced as contain histones that are hypoacetylated and methylated. Thus, we prepared a chip seeded with NHEKs in a chamber and Vero E6 cells in the other as “sensor” cells. Then keratinocytes were infected with high concentrated recombinant virus (2.5 × 10^2^ PFU/mL). ACV was added following two different timing: pre and post-infection, or only after virus adsorption. Next, supernatants were collected at different time points and finally assayed for infectious virus on Vero E6 cells seeded on plates (**Figure S3A**). Infectious virus was detectable in the supernatant of infected NHEK cultures, but it was never detected in the supernatant of the uninfected chamber. This validates our microfluidic model for the study of the neuron-to-cell virus spreading.

### Evaluation of the HSV-1 lytic infection in keratinocytes and neurons

The main cellular components for HSV tropism are represented by epithelial cells and sensory neurons. To obtain the most physiological and reproducible infection *in vitro*, we decided to evaluate the response to HSV of different cell types, including primary and immortalized keratinocytes, as well as immortalized and DRGO neurons.

The first step was to assess if all the different cell types to be used were permissive to the lytic infection of this virus strain. Thus, both primary (NHEK) and immortalized (HaCaT) human keratinocytes were infected, as well as neuroblastoma cells (SH-SY5Y) and 3D cultured human iPSC-derived DRGOs. Epithelial cells (Vero E6) were used as a positive control of infection. After 24 hours, the expression of ICP0 and gC was investigated by live imaging on infected cells, and all the tested cell types showed the effects of productive and lytic infection (**Figure S4**). Interestingly, the immunofluorescence signal reflected the different timing of HSV-1 gene expression during productive infection with the EGFP signal (ICP0 promoter, immediate early gene) significantly more widespread than the RFP one (glycoprotein C promoter, early gene). These results confirm that, similarly to other cellular models, DRGOs can be efficiently infected with HSV-1, establishing a lytic infection.

### Successful establishment of HSV-1 latency in DRGO neurons

As virus reactivation from latently infected cells was a main objective of this study, a novel protocol for HSV-1 establishment of latency was set up using two virus strains: HSV-1 HF, a laboratory strain, and the recombinant HSV-1 KOS. The induction of HSV latency is performed by pre-treating cells with Aciclovir (ACV) before virus adsorption. Therefore, a fundamental requirement for both HF and KOS viral strains is their susceptibility to ACV. To this aim, *in vitro* phenotypic assays were performed, and results confirmed that the amount of antiviral to be used in the latency protocol would be well above their IC50 (**Figure 4A**).

**Figure 4.**
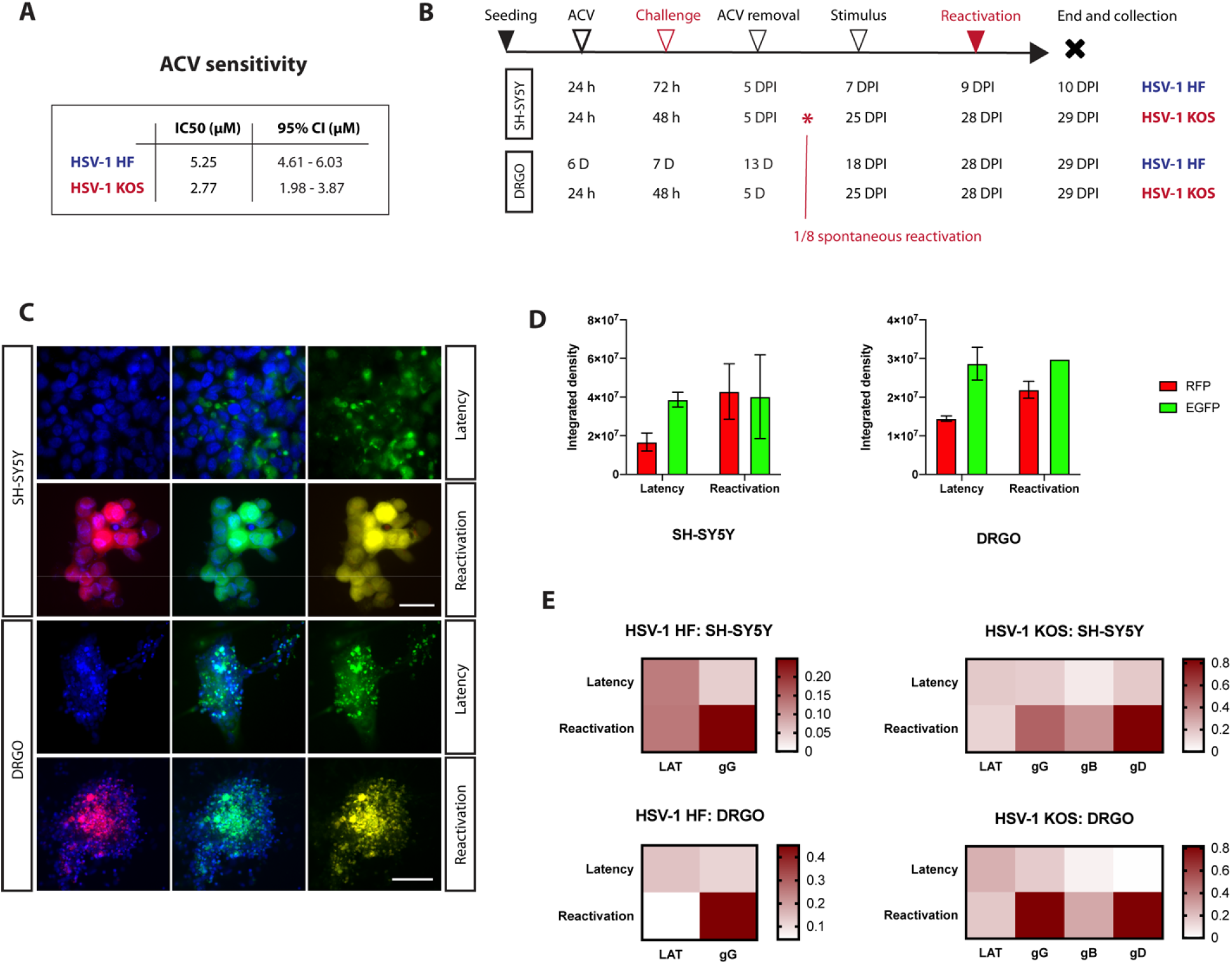
HSV-1 latency and reactivation protocol setup. **(A)** ACV phenotypic assay performed on HSV-1 laboratory strain (HF) and recombinant virus (KOS). **(B)** Schematic representation of different protocol timing tested on SH-SY5Y and DRGOs, using both virus strains. **(C)** Immunofluorescence analysis SH-SY5Y and DRGOs during virus latency and reactivation. Virus protein gC (red, first column) is transcribed only during productive infection, while ICP0 promoter (green, second column) is active during both stages of infection. Total nuclear DNA is counterstained with Hoechst (blue), the last column shows the merge of green and red signals. Scale bars, SH-SY5Y 30 μm, DRGO 200 μm. **(D)** Levels of ICP0 (green fluorescent signal) and gC (red fluorescent signal) expression measured as integrated density values for SH-SY5Y and DRGOs during latency and reactivation. Mean ± SD is reported. **(E)** Heatmap showing virus gene expression analysis of SH-SY5Y and DRGOs during latency and reactivation using both HSV-1 strains (dark red= high expression; white = no expression). Each condition was tested in quadruplicate.

The HF non-recombinant strain was used to tune our experiments as latency protocol on SH-SY5Y was already described in literature (Sun et al. 2016). SH-SY5Y cells were infected with different virus concentrations after ACV pretreatment (**Figure 4B**). After adsorption, cells were maintained in culture media supplemented with ACV for 5 days, and then cultured without antiviral for 2 days to monitor for spontaneous reactivations. Then, thermal stress was used for inducing virus reactivation, and the cytopathic effect caused by productive infection was visible 2 days later. Latency establishment and virus reactivation were obtained by infecting with 4 PFU/mL of the HSV-1 HF strain. Next, we adapted and validated a similar protocol for DRGO-derived neurons. Reactivation stimulus was given 5 days following ACV trigger, and about 10 days later the cytopathic effect on cells started to appear. Noteworthy, virus reactivation from latency was successfully obtained even using three-dimensional cultures. Once the protocol was developed using a reference strain, the recombinant KOS virus was used. First test on SH-SY5Y cells confirmed both the optimal virus concentration to infect cells without immediate lysis of the monolayer (2.5 PFU/mL), and the appropriate timing: 20 days between antiviral removal and stimulus to obtain detectable reactivation 3 days later. This dilation time after ACV removal from the medium was mandatory to ensure that what we observed was indeed a controlled and not a spontaneous reactivation (Danaher, Jacob, Chorak, et al. 1999). With this procedure, we observed spontaneous reactivation in 1 out of 8 total wells in 11 DPI. Then, DRGOs were latently infected and reactivated using the same experimental protocol. Interestingly, no spontaneous reactivation occurred at that time.

Live imaging was obtained of latently infected cells in culture, as the gene encoding the ICP0 protein of HSV-1 is located in the region of the HSV-1 genome encoding LATs (Block & Hill 1997) and it is transcribed during latency (S.-H. Chen et al. 2002). Indeed, it was possible to detect the presence of latently infected cells before stimulus in both SH-SY5Y and DRGOs cultures as highlighted by the EGFP signal (**Figure 4C**). Lytic infection could be visible only after thermal stress, monitored by the combination of both GFP and RFP fluorescence signals. Integrated density was used to compare both green and red fluorescent signals of reactivated and latently infected cells, highlighting the actual difference only between the RFP signal detected in both SH-SY5Y and DRGOs, even if without statistical significance (**Figure 4D**).

To assess the differences in gene expression profiles during latency and reactivation, cells were harvested at the end of the experimental protocol for total RNA extraction. Quantitative Two-step Reverse transcription-PCR (RT-qPCR) showed a slight but no significant reduction of LAT transcripts 24h post-reactivation in both SH-SY5Y cells and DRGOs, using both viruses (**Figure 4E**). Instead, an important increase of gG expression was observed in both HSV-1 HF and KOS infected cells (p<0.0001 and p<0.01, respectively) and DRGOs (p<0.0001 using both viruses). Next, we further characterized the recombinant virus infection analyzing also other early and late gene expression. We observed an increase in the expression of both glycoprotein D (gD) and B (gB), although the latter was higher up-regulated (p<0.0001) than the former (p<0.05 only in SH-SY5Y). It is interesting to note that during the latent phase, SH-SY5Y cells retain partial expression of lytic products, as well as lytic RFP fluorescence, while in DRGOs is almost absent, suggesting a more physiological and sustainable viral latent phase in the human organoid model. In summary, all data confirmed that we were able to establish stable and long-term HSV-1 latency into DRGOs, allowing us to better model virus reactivation in human sensory neurons using three-dimensional cultures directly differentiated from hiPSCs.

### Testing HSV-1 latency in the microfluidic culture system

Given the results described above, two experimental settings were tested for the study of HSV-1 reactivation from latency, the “One-way” experiment (EXP #1) involving direct infection and latency of DRGOs (**Figure 5A, B**), and the “Back and forth” protocol (EXP #2) with establishment of latency into DRGOs following NHEK infection (**Figure 5C, D**), summarizing the physiological behavior of HSV-1. For both experiments, NHEKs were seeded together with DRGOs to allow axonal contacts, and again 18 days later to fill the chamber. After 21 days all chambers were pretreated with ACV and infected with recombinant HSV-1 KOS, except for NHEK of EXP #2, which were treated only 6 h post adsorption to allow virus replication and spreading to DRGOs through axonal connections and receptor recognition. All cells were cultured with proper media supplemented with antiviral for 5 days. Then, it was removed, and thermal stress was used for inducing virus reactivation. The cytopathic effect caused by productive infection was visible 2 days later. Supernatants from different time points were tested on Vero E6 monolayers. While infectious virions were detectable in the supernatant of NHEK after virus adsorption (EXP #2), it was never detected in the supernatant of uninfected cultures, confirming that virions never passively transfer between chambers (**Figure S3B**), making our neurosensory-epithelia connection on-a-chip a feasible tool to model HSV-1 reactivation processes *in vitro*.

**Figure 5.**
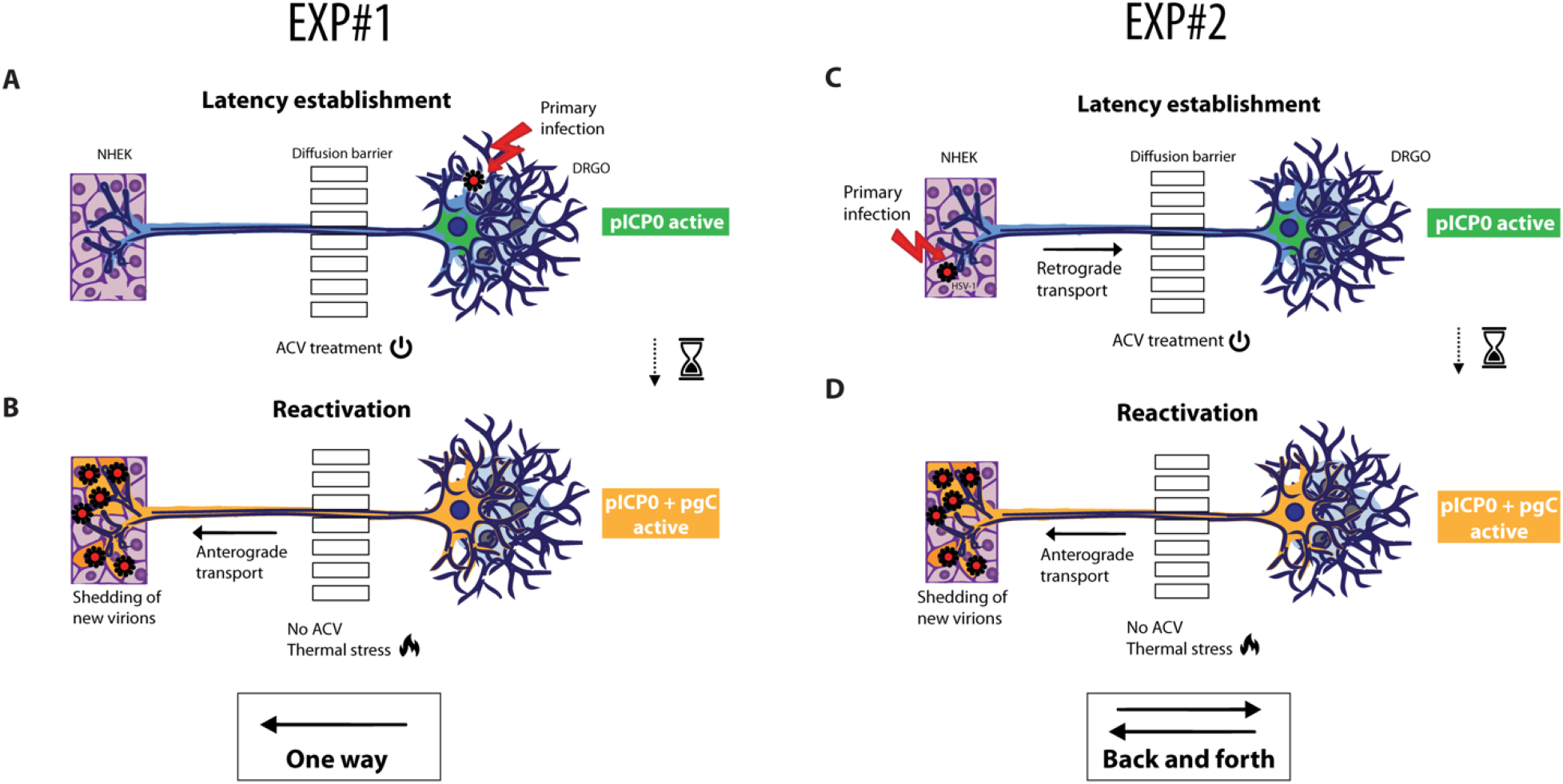
Testing HSV-1 latency in the microfluidic culture system. Graphical representation for HSV-1 latency establishment directly in DRGOs following one-way reactivation protocol ( EXP#1, **A-B)** or back and forth experiment (EXP#2, **C-D)**. In EXP#1, organoids were infected **(A)** and latency obtained through ACV addition to culture medium. Thermal stress **(B)** led to controlled reactivation, and anterograde transport of virus particle resulted in NHEK lytic infection. EXP#2 instead was carried out by HSV-1 latency establishment indirectly in DRGOs **(C)**. NHEK were infected, and retrograde transport of virus particles resulted in HSV-1 latency establishment in organoids. Then, thermal stress **(D)** led to controlled virus reactivation, and anterograde transport of virus particles resulted in NHEK lytic infection.

### Setting the “One-way” viral infection in the neuro-epithelial culture system

The first experiment involved direct infection of DRGOs with low titer HSV-1, and establishment of latency by ACV supplemented medium **(Figure 5A)**. Gene expression profiles were assessed, and exclusively LAT transcripts were detected in DRGOs only, confirming once again both latency protocol efficacy and no passive virus diffusion occurring through the NHEK chamber **(Figure 6A)**. When controlled reactivation occurred following ACV removal and thermal stress, the virus spread to NHEKs through axonal connections to keratinocytes in a “one way” model of HSV-1 infection from reactivation **(Figure 5B)**. Analysis of gene expression revealed that LAT transcripts were not significantly decreased 24h post-reactivation in DRGOs, as already found in mono-culture data **(Figure 6B)**. They were different in NHEK (p<0.01), as no transcripts were detected in that chamber during latency. No early (gG and gB) and late (gD) genes expression was detected during latency in both DRGOs and NHEK, but they were actively transcribed during reactivation in both cell chambers, as shown by graphs (DRGOs: gG p<0.001; gB p<0.0001; gD p<0.0001. NHEK: gG p<0.0001; gB p<0.05; gD p<0.0001).

**Figure 6.**
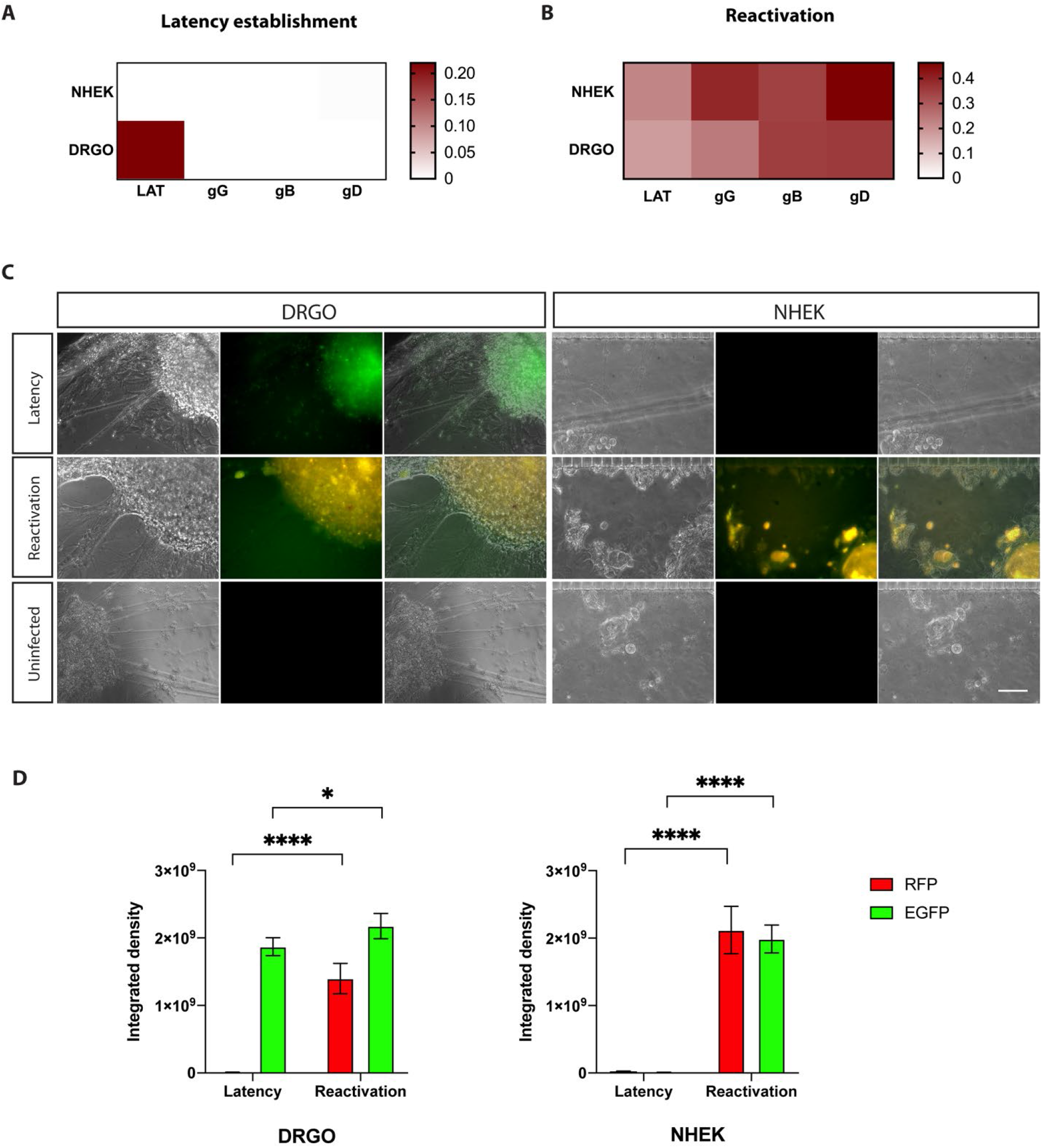
One-way experiment (EXP#1). Virus gene expression profiling for HSV-1 latency establishment **(A)** directly in DRGOs. Organoids were infected and latency obtained through ACV addition to culture medium. Thermal stress led to controlled reactivation **(B)**, and anterograde transport of virus particle resulted in NHEK lytic infection. **(C)** Live imaging of DRGOs and NHEK showing recombinant HSV-1 during latency (pICP0 active, green) and reactivation (pICP0 and pgC active, yellow). Scale bar, 100 μm. **(D)** Levels of ICP0 (green fluorescent signal) and gC (red fluorescent signal) expression measured as integrated density values for DRGOs and NHEK, during latency and reactivation. Mean ± SD, *p<0.05; ****p<0.0001, n = 6 independent experiments (12 DRGOs).

Live imaging of the two cell chambers was concordant with gene transcripts analysis (**Figure 6C**). Latency was detected by the EGFP signal only in DRGOs. After the induction stimulus, the productive infection was detected in both the neuron and keratinocyte chambers thanks to the co-presence of EGFP and RFP fluorescence. Values of fluorescent intensity confirmed what observed: lytic red signal was detected only during productive infection in both cell populations (p<0.0001), while green signal was present in NHEK images only after the induction stimuli (p<0.0001). DRGOs showed EGFP fluorescence in both latent and reactivate states, albeit with a slight but significant difference (p<0.05) (**Figure 6D**). No signal was detected in uninfected microfluidic chips, confirming that what observed was effectively due to the virus reactivation process.

### Validation of the “back and forth” viral spreading

We, then, conceived EXP #2 which is more complex since it is designed to test the “back and forth” viral spreading. For this, we defined a cycle starting with the infection of keratinocytes, virus migration to neurons for latency establishment and reactivation followed by virus migration back to keratinocytes which closely models the kinetics and topography of the natural infection. To establish this model in microfluidics, NHEKs were directly infected, and ACV was applied 6 h later to allow a minimum of virus replication and spreading through axons before latency establishment into DRGOs (**Figure 5C**). As neurons are the preferred cell type in which to recapitulate latent infection in cell culture, it was already observed that latent state cannot be detected in non-neuronal cells (Thellman & Triezenberg 2017). That was further validated through gene expression profiles, as no other virus transcript rather than LATs were detected in DRGOs and none in keratinocytes **(Figure 7A)**. Then, ACV removal and thermal stress drove HSV-1 reactivation and neuron-to-cell spreading, leading to active replication in keratinocytes (**Figure 5D**). Once again LAT transcripts were not significantly decreased 24h post-reactivation in DRGOs, while they were actively transcribed in NHEKs only after reactivation (p<0.0001) **(Figure 7B)**. Early (gG and gB) and late (gD) genes were actively transcribed only during reactivation in both chip compartments, and overall lower levels of expression were observed in EXP #2 than #1 for both DRGOs and NHEKs, as shown by qPCR analysis (DRGOs: gG p<0.0001; gB p<0.01; gD p<0.001. NHEK: gG p<0.0001; gB p<0.001; gD p<0.0001).

**Figure 7.**
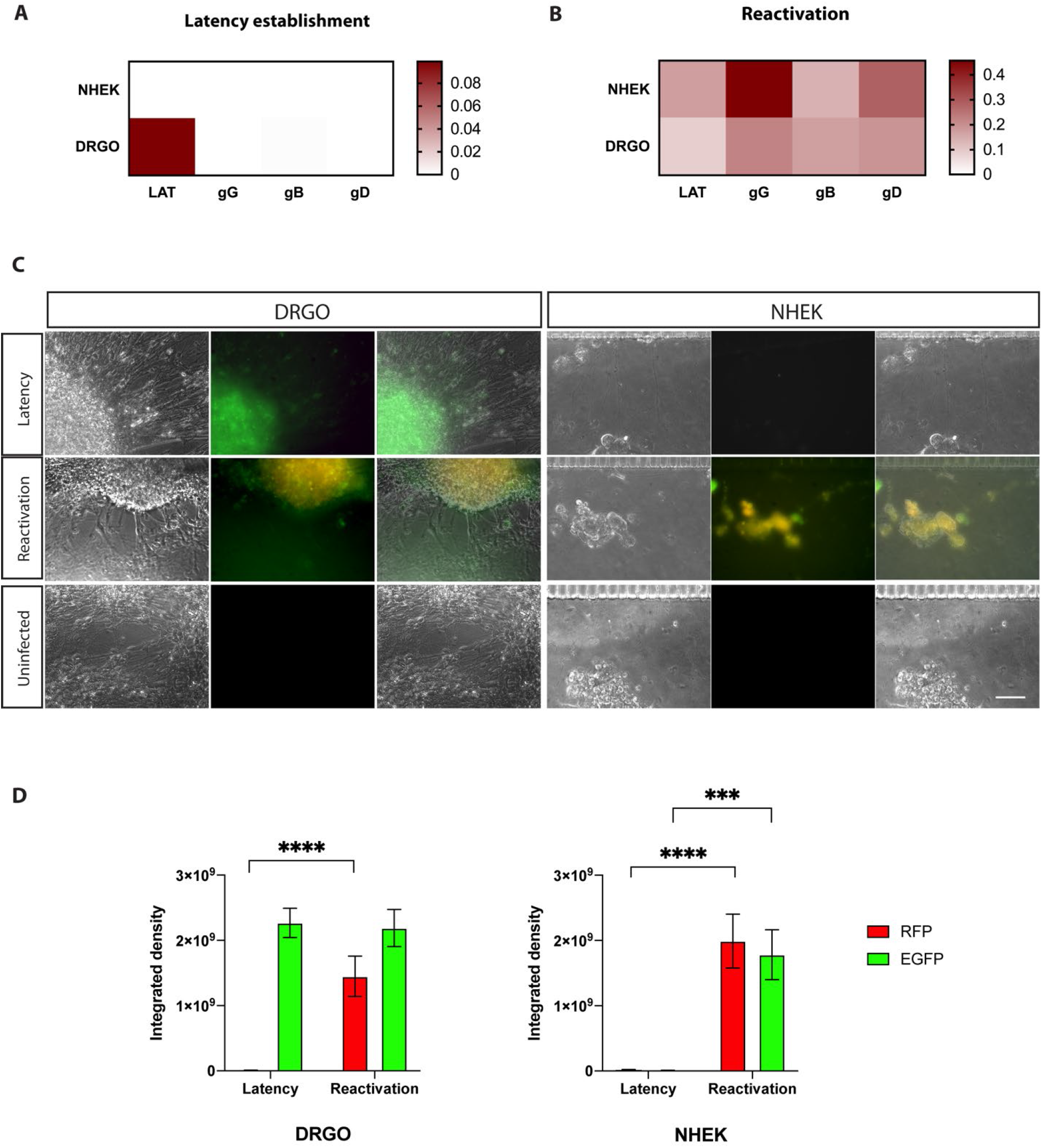
Back and forth experiment (EXP#2). Virus gene expression profiling for HSV-1 latency establishment **(A)** indirectly in DRGOs. NHEK were infected, and retrograde transport of virus particles resulted in HSV-1 latency establishment in organoids. Thermal stress led to controlled reactivation **(B)**, and anterograde transport of virus particle resulted in NHEK lytic infection. **(C)** Live imaging of DRGOs and NHEK showing recombinant HSV-1 during latency (pICP0 active, green) and reactivation (pICP0 and pgC active, yellow). Scale bar, 100 μm. **(D)** Levels of ICP0 (green fluorescent signal) and gC (red fluorescent signal) expression measured as integrated density values for DRGOs and NHEK, during latency and reactivation. Mean ± SD, ***p<0.001; ****p<0.0001, n = 6 independent experiments (12 DRGOs).

This conclusion was further confirmed by living cell imaging (**Figure 7C**). In fact, EGFP signal was detected only in DRGOs, corroborating latency establishment exclusively into neurons. As expected, fluorescence was less intense than EXP #1 due to the indirect modality of neuronal infection. After the induction stimulus, the productive infection was detected in both cell chambers. These data are confirmed by immunofluorescence signals measurements: RFP signal was detected only during productive infection in both cell populations as for EXP#1 (p<0.0001), and EGFP signal in NHEK images only after the induction stimuli, but less intensely than in the previous experiment (p<0.001). DRGOs showed an important green signal as well in both latent and reactivate states, but this time without significant increase (**Figure 7D**).

In this case as well, no fluorescence signal was detected in microfluidic chips with uninfected cells, validating live imaging observations. These results showed that DRGOs were successfully infected by HSV-1 through retrograde axonal transport from infected keratinocytes, and stress stimulus led to direct neuron-to-cell spreading of infection. Microfluidic chips were maintained in the latent state as control of spontaneous reactivation until the end of the experiment (8 DPI). They were not subjected to thermal stress after ACV removal from the culture medium, and cells were collected to test gene expression. As expected, only LAT transcripts were detected in DRGOs, while no virus gene expression was detected in keratinocytes (**Figure S5**).

Collectively the presented data validate the establishment of a new human sensory neuron-to-keratinocyte circuit on-a-chip suitable to fully recapitulate the key HSV-1 viral processes of reactivation from latency.

## Discussion

Studies on neurotropic viral reactivation are hampered by the lack of a suitable experimental system of human origin that recapitulates the cellular complexity and stereotyped connectivity. From a therapeutic point of view, only systems that are fully human in their composition allow a reliable characterization of drug efficacy and identification of novel targets. In this regard, several cellular settings have been tested in co-culture, using either human and non-human-derived primary or immortalized cells, but none has enabled to reconstitute both latency and reactivation dynamics of neurotropic viruses trough a neuron-to-cell functional connection to date (Sun et al. 2016; Markus et al. 2015; Pourchet et al. 2017; D’Aiuto et al. 2019). The direct consequence of this is the absence of a fully human-based model where to faithfully test drug candidates. In fact, at the moment it is impossible to draw a general prediction for the overall activity of novel drug compounds or checking the potentials of antivirals to be included in clinical trials using a single experimental system. Moreover, the lack of a human neuron-skin circuitry model hampers the possible identification of novel correlates of protection from infection, including the cell-to-cell spread, and the diagnosis of drug resistance (Criscuolo et al. 2019). At present, the preclinical development of novel effective antiherpetic compounds is mainly based on cell culture studies and has been limited in the last decades to improvements of nucleoside analogs. In other words, no novel antiherpetic drug classes for systemic use reached the clinic (Clementi et al. 2016). Other steps of the virus life cycle have been addressed by both novel preclinical compounds (Clementi et al. 2017; Baram-Pinto et al. 2010; Mohammed Fayaz et al. 2012; Krawczyk et al. 2013; Luganini et al. 2011; Bergaoui et al. 2013; Price et al. 2011; Criscuolo et al. 2018) and vaccine strategies (Veselenak et al. 2012; Stanberry et al. 2002; Casanova et al. 2002), but no one has proven to be effective to date or, if so, has already reached clinical development.

Here, we described and validated a novel *in vitro* human cellular system on-a-chip that could entirely fill this current gap. The human neuronal component is relying on the use of DRGOs, DRG-like organoids derived by 3D differentiation of hiPSCs into the plethora of sensory neurons and glial cells, and capable of directly connecting with keratinocytes. These findings confirm and further extend the intrinsic capacity of DRGO sensory neurons to contact and make stable connections *in vitro* with the authentic peripheral target tissues of DRGs in human body, as we initially showed for the reconstitution of the muscle spindles together with the intrafusal muscle fibers (Mazzara et al. 2020). Thus, this human neuron-skin circuit on-a-chip represents an invaluable platform to study viral mechanisms and kinetics in a patterned circuit fundamentally similar to its *in vivo* counterpart and enable biochemical and molecular analyses that would be difficult, if not impossible, to perform *in vivo*.

The establishment of this system has required numerous steps of optimization. At first, we observed that co-cultures of DRGOs and NHEKs were possible only if the microfluidic chip was used to prevent medium mixing, as both of them resulted in a detrimental effect for one of the two cell populations. We hypothesized that the serum present in the HaCaT medium could be the possible cause of this effect on DRGO-neurons. Free axonal endings developed in the co-cultures on the chip were characterized only when primary keratinocytes (NHEK) were used, but not immortalized cells (HaCaT), probably, again, for the presence of the serum that impaired axonal growth and subsequent connections to keratinocytes. Finally, no passive diffusion of infectious virus particles was detected between chambers in the chip, even using 2 logs more concentrated virus than used in latency protocol. Furthermore, Vero E6 cells seeded into the opposite chamber as sensor cells because of their high permissiveness to infection remained uninfected, demonstrating the reliability of these devices for studying the axonal passage of the virus.

Then, the possibility of having a pathogen permissive neuronal model able to be correctly recognized from the virus, confirmed the feasibility of DRGOs as a suitable model for *in vitro* studies on human genetic and infective disease affecting peripheral sensory neurons, and allowed us to set-up neuron-to-skin connectivity, suitable for multiple studies, including HSV-1 tropism. We showed that the pathogen competence of DRGOs is due to the presence of many specific HSV receptors, confirmed by global transcriptome profile. DRGOs showed similar expression to what was observed in primary ganglia for the principal HSV receptor *NECTIN1*, but also *TNFRSF14* (*HVEM*), PILRA, *NRP1*, *MYH9*, *ITGA5*, *ITGA8*, *ITGB6* and *ITGB 8*, as well as the expression of cardinal genes for the heparan sulfate biosynthesis and integrins. Noteworthy, through mining of scRNA-seq datasets, we revealed that DRGO cell types express a unique combination of HSV receptors. These surprising findings might suggest that kinetics and efficacy of HSV infection is not identical among the different DRG neuronal populations providing a different impact on the overall viral pathological effects. Furthermore, it would be interesting to understand the transcriptional cascade which links a particular pattern of expressed viral receptors with the specific identity of the sensory neuronal cell type.

In this study we showed that the primary infection of human DRGOs and primary adult human NHEKs with the HSV-1 engineered strain resembled that described for the reference epithelial immortalized cell lines (Vero E6, HaCaT) and neuroblastoma-derived cell line (SH-SY5Y). This demonstrates that the engineered HSV-1 strain retains the broad tropism of the wild type virus. Subsequently, the setup of an effective latency and reactivation protocol was further established. HSV-1 laboratory strain (HF) and SH-SY5Y were initially used to help in adapting previous protocols to our specific conditions. Primary infection using low titer virus and cell medium supplemented with ACV led to latency establishment into cells, and thermal stress led subsequently to controlled reactivation of infected cell culture, as performed with other HSV-1 strains. Gene expression profiles were assessed to confirm the two stages of infection, analyzing LAT and an early gene (gG) transcription. Thirdly, recombinant HSV-1 was used but the time between ACV removal and stimulus was lengthened to monitor possible spontaneous reactivations. Indeed, despite spontaneous reactivation was occurred in one out of eight replicates, the protocol has substantially proven to be reliable also for this virus strain. Finally, gene expression analysis was performed and gB (early gene) and gD (late gene) transcripts were investigated. All of the results confirmed that both latency establishment and controlled reactivation were possible using the recombinant HSV-1. This greatly facilitated the subsequent experimental procedures, as the recombinant virus allowed live imaging discrimination of cells in a latent state or reactivated. Its reporter genes (EGFP and RFP) are under the control of the latency viral promoter ICP0 and lytic viral promoter glycoprotein C, respectively. Therefore, latently infected cells are easily identified for the green fluorescent signal, which turns yellow if there is a lytic infection following reactivation. Latency protocol was assessed on DRGOs using both laboratory and recombinant virus strains. The results were even more convincing as no spontaneous reactivation occurred, as confirmed by live imaging and gene expression analysis.

Our system provides a reliable model where to study HSV-1 reactivation from latency into neurons and subsequently infection of keratinocytes through neuron-to-cell connectivity. The first on-a-chip experiment we performed was a direct infection of DRGOs with low titer recombinant HSV-1 and in the presence of ACV to obtain latency establishment. Both cell live imaging and gene expression profiling confirmed the achievement of this result, as only green fluorescence and LAT transcripts were detected in DRGOs. Conversely, NHEKs remained virus-free as demonstrated by the lack of passive diffusion of infective virus particles. Then thermal stress led to virus reactivation into neurons, and immunofluorescence and gene expression analysis demonstrated productive infection in both chip chambers. Confirming that no passive diffusion is possible within microfluidic compartments, and HSV-1 reactivation from latently infected DRGOs led to NHEK primary infection only through axonal contacts. As already observed in mono-culture results and literature data (Sun et al. 2016; Danaher, Jacob, Chorak, et al. 1999), 24 hours from reactivation were not sufficient to detect a significant decrease of LAT transcripts but were enough to observe a significant expression of early genes (gG and gB) and the late HSV gene gD. The differences observed between NHEKs and DRGOs are in agreement with the increased susceptibility of epithelial cells to productive infection compared to neurons (Kent & Fraser 2005; Thellman & Triezenberg 2017).

The positive results obtained with this first experiment allowed us to set a system to better model the physio-pathological conditions of HSV-1 post-infection reactivation from latency occurring *in vivo*. In this experimental setting, NHEKs were primarily infected without ACV to facilitate virus replication and spreading through retrograde transport to DRGOs. Conversely, subsequent supplementation with ACV blocked virus productive infection into the keratinocytes concomitantly allowing the establishment of latency into neurons as confirmed by both cell live imaging and virus transcripts analysis. The lower intensity compared to the EXP #1 conditions, is probably due to the greater complexity of the model that resulted in fewer DRGO neurons infected due to retrograde migration compared to direct infection. In fact, in the first experiment, the virus adsorbed on the whole organoid whereas in the retrograde infection the latency is established only by the portion of viruses undergoing retrograde trans-axonal migration. No lytic or latent infection was detected in the NHEK chamber. Supernatants tested on sensor cells corroborated the presence of infective virus particles only into the NHEK chamber after adsorption, but neither into the DRGO compartment nor in both compartments in the subsequent time points. Thus, we successfully obtained latency into DRGO neurons from HSV-1 replication into keratinocytes and the subsequent virus spreading through axonal contacts and retrograde transport, closing our “back and forth” reactivation cycle in the microfluidic platform. Interestingly, transcriptional analysis from NHEKs showed an imbalance towards early gene expression, suggesting a possible lower virus titer able to reach keratinocytes or a delayed timing of anterograde transport compared to what resulted from the simplified method described above.

These results confirmed that our neuron-skin connectivity is a reliable platform to study at molecular level HSV axonal transmission during reactivation from latency in a topographic co-culture system. Its fully human composition and the miniaturization and standardization, made possible by microfluidics technology, provide this device with unique features to date, and perfectly fitted for several applications, spanning from the fine dissection of molecular mechanisms occurring during neurotropic virus infection to the translational application for both therapeutic and diagnostic purposes.

The development of stable neuron-to-keratinocyte connectivity into a microfluidic device enables the characterization of molecular mechanisms involved in the reactivations and spreading of neurotropic viruses in a fully human context. Next, our new knowledge on the viral receptors expressed by human neurons will facilitate the mechanistic dissection of the complex virus-host interactions. This on-chip platform has also important application for the validation of HSV infection biomarkers, treatment responses, or potential curative therapeutic interventions, which cannot be studied using other *in vitro* or even *in vivo* systems. Furthermore, our new model was validated by using a well-characterized recombinant virus. Therefore, new studies using also clinical virus isolates will be important for shedding light on unknown pathophysiological aspects of herpesvirus reactivation. Finally, the generation of organoids with defined cellular populations will enable the development of drug screening platforms. This opens the door to further expanding this culture platform to other neurotropic viruses to dissect reactivation mechanisms and to identify novel targets for antiviral therapy.

## Acknowledgments

Experiments employing the Leica confocal SP2 microscope, and Zeiss Axio Observer.Z1 microscope with QImaging Exi-Blue were carried out in ALEMBIC, an advanced microscopy laboratory facility established by the San Raffaele Scientific Institute and the Vita-Salute San Raffaele University. This work was partially supported by the European Research Council to V.B. (AdERC #340527; ERC-PoC #842423).

## Author contributions

PGM, EC, NC, VB, MC conceived and designed the study, PGM and EC performed all the experiments, analyzed all the data, assembled the figures and wrote the manuscript. MR and CP generated the microfluidic chips. MC and NM provided advice and contributed to data analysis and manuscript preparation. NC, VB, MC and RB secured founding and supervised the study. All authors read and approved the final manuscript.

## Methods

### Cell lines

Vero E6 (Vero C1008, clone E6 – CRL-1586; ATCC), SH-SY5Y (CRL-2266; ATCC) and HaCaT (kindly provided by Cremona O., (Boukamp et al. 1988)) cells were cultured in Dulbecco’s Modified Eagle Medium (DMEM) supplemented with non-essential amino acids (NEAA), penicillin/streptomycin (P/S), Hepes buffer and 10% (v/v) Fetal bovine serum (FBS). Primary NHEK cells from singe adult donor (C-12005; PromoCell) were cultured in Keratinocytes Growth Medium 2 (C-20011; PromoCell). Human iPSCs from healthy control fibroblasts (DIGI and Neof2, obtained from the IRCCS Carlo Besta Neurological Institute and ATCC, respectively) where reprogrammed using the CytoTune-iPS 2.0 Sendai Reprogramming Kit (A16517; Thermo Fisher Scientific), maintained in feeder-free conditions in a mTeSR™1 (85850; Stem Cell Technologies) medium supplemented with 1% Penicillin-Streptomycin (P0781; Merck) in 6-well culture plates coated with Matrigel^®^ hESC-Qualified Matrix (354277; Corning).

### Antiviral compound

Aciclovir (ACV; 9-[(2-hydroxyethoxymethyl) guanine]) (A-4669; Merck) was dissolved in DMSO at a concentration of 10 mg/mL stored as single-use aliquots at −20 °C. Dilutions were made in cell medium immediately before use.

### Generation of 3D DRG-like organoids (DRGOs)

iPSC-derived dorsal root ganglia organoids (DRGOs) were developed as follows. 9×10^3^ iPSCs are seeded into low-adhesion V-bottom 96-multiwell plates (277143; Thermo Fisher Scientific). The following day change medium with KSR medium (DMEM-F12 with 15% KSR, 1% of each: pen/strep; Non-Essential Amino Acids, and β-mercaptoethanol 100 μM and L-Glutamate 2nM). The day after (DIV 0) add KSR medium plus SB431542 10 μM (S4317; Merck) and LDN193189 100 μM (04-0074; Stemgent). The medium is subsequently changed every 48hrs. Between DIV 4 and DIV 9, CHIR99021 3 μM (130-103-926; Miltenyi); SU5402 3 μM (SML0443; Merck); and DAPT 10 μM (D5942; Merck) are added and KSR medium is gradually switched to N2 medium (Neurobasal medium (21103049; Thermo Fisher Scientific); N2 (17502-048; Thermo Fisher Scientific); pen/strep; NEAA, and L-Glutamine 2mM). On DIV 10 add 100% N2 medium plus recombinant human Brain Derived Neurotrophic Factor 10ng/ml (BDNF 450-02; Peprotech); Glial-Derived Neurotrophic Factor 10ng/ml (GDNF 450-10; Peprotech); Nerve Growth factor 10ng/ml (NGF N6009; Merck); Neurotrophin-3 10ng/ml (NT-3 450-03; Peprotech) and Ascorbic Acid 200 μM (AA 49752; Merck). On Day 16, organoids were plated into Matrigel®-coated 24-well plates with and without glass coverside, directly onto cultures of Keratinocyte or in microfluidic devices and maintained in N2 medium supplemented with the maturation factors or in different combinations of mediums where specified. Replacing half of the appropriate medium every 72 hours until samples are collected for analysis or for up to 90 days. Two fluorodeoxyuridine (FUDR) treatments were performed from DIV 17 to 26.

### Microfluidic culture system

A microfluidic compartmentalized bioreactor was designed, featuring two symmetric lateral chambers (43 μm high, 1.5 mm wide and 7 mm long), one for the neuronal culture and the other one for the epidermal culture. Each compartment is connected to two wells at its ends, serving as culture medium reservoirs during cell culture. Additionally, the neuronal compartment is provided with two smaller reservoirs dedicated to host the soma of DRGOs. The two chambers are fluidically connected by 250 microgrooves, with a width of 5 μm, a height of 6 μm and a length ranging from 830 to 930 μm. Microgrooves were conceived to guide axon growth, while preventing convective flow between compartments.

Microfluidic devices were fabricated through soft lithography of PDMS on master molds. Briefly, the chamber design except for the microgrooves, was printed on a high-resolution photomask (64’000 dpi). The master molds were fabricated in a cleanroom environment through a two-step photolithography procedure of SU-8 (Microchem) on silicon wafers. First, the microgrooves pattern was created by direct laser writing of a layer of SU-8 2005 with a thickness of 6 μm, by using a mask-less laser writer (MLA100, Heidelberg). Subsequently, a 43 μm thick-layer of SU-8 2035 was spin-coated onto the wafer and exposed to a collimated UV beam through the photomask containing the chambers features. The PDMS layers were then fabricated by replica molding. PDMS (SYLGARD^®^ 184 Silicone Elastomer Kit, Dow) was poured on silicon wafers at a pre-polymer to curing agent mixing ratio of 10:1 (w/w) and cured at 65°C for 3h. After the curing phase, PDMS was peeled-off the molds, trimmed and through-holes were punched to obtain the lateral reservoirs (8 mm diameter) and the DRGOs seeding sites (2 mm diameter). Finally, after plasma activation (Harrick Plasma), the PDMS layers were bonded to 24×50 mm microscope glass slides (Superslip^®^ Cover Glass, TED PELLA).

The compartments were coated with Matrigel^®^ 1:200 (Thermo Fisher Scientific) and incubated overnight at 4°C. Each compartment holds about 150 μL of medium. Two 16 days old DRGOs were seeded into the dedicated seeding sites, 5 × 10^4^ NHEK cells were seeded in the other compartment (**Fig. 1A**). Neuron and keratinocytes cultures were maintained as described above.

### Transcriptomic analysis

Bulk RNA-seq fastq files were mapped to the hg38 mouse reference genome with the Bowtie2 aligner. Differential gene expression and functional enrichment analyses were performed with DESeq2 (Cagnoli et al. 2004) and GSEA (Rubio et al. 2016), respectively. Statistical and downstream bioinformatics analyses were performed within the R environment.

Gene expression heatmaps were produced with GENE-E (Broad Institute). Data were deposited in the NCBI Gene Expression Omnibus repository with the GSE133755 GEO ID.

RNA-seq datasets for human somatic dorsal root ganglia (SRA SRP077657) (Chambers et al. 2009) and human iPSCs (GEO GSE120081) were obtained from the NCBI GEO/SRA repositories for data mining.

Gene ontology aggregated categories were produced by combining multiple GO datasets: *Heparan sulfate biosynthetic pathways* (GO Heparan Sulphate Proteoglycan Binding; GO Heparan Sulphate Proteoglycan Biosynthetic Process, GO Heparan Sulphate Proteoglycan Biosynthetic Process Enzymatic Modification, GO Heparan Sulphate Proteoglycan Biosynthetic Process Polysaccharide chain B Biosynthetic process, GO Heparan Sulphate Proteoglycan Metabolic Process), Integrins signaling genes (GO Integrin Mediated Signaling Pathway).

ScRNA-seq fastq files were aligned to the GRCh38 human reference genome with cellranger count (10x Genomics) with default parameters, and human Gencode v33 annotations (PMID:30357393). Gene count matrix log-normalization (10,000 scale factor), gene clustering, dimension reduction analysis (UMAP), differential gene expression analysis, and plotting were performed with Seurat (Stuart et al. 2019) within the R environment (R Core Team (2020). R: A language and environment for statistical computing. R Foundation for Statistical Computing, Vienna, Austria. URL: https://www.R-project.org/). Functional enrichment of gene ontology biological processes datasets was performed with DAVID (Huang et al. 2007).

### Viral infections

The laboratory strain HSV-1 HF (VR-260; ATCC) and a recombinant fluorescent virus kindly provided by Dr. P. Kinchington from University of Pittsburgh were used. It is a KOS-based (VR-1493; ATCC) recombinant strain incorporating enhanced green fluorescent protein (EGFP) and monomeric red fluorescent protein (RFP) as reporters whose gene expression is driven by the viral promoters ICP0 and glycoprotein C, respectively (Decman et al. 2005). Lat is the only actively transcribed gene from the virus in latently infected cells, and the boundary between transcription-permissive and -nonpermissive regions is located near the 3′ end of the ICP0 (Amelio et al. 2006). Thus, the ICP0 promoter is actively transcribed during latency, whereas the surrounding regions, such as the lytic-specific gene ICP0, are silenced.

### Infection of 2D cultures

For lytic infections, cell-free virus was adsorbed onto monolayer of Vero E6, HaCaT, NHEK or SH-SY5Y seeded on Matrigel^®^ coated slides with a removable 12 well silicone Chamber (Ibidi). After 2 h the cells were washed and cultured for 24 h, when live images were acquired. For latent infection, SH-SY5Y cells were preincubated with ACV 100 μM in complete DMEM supplemented with 2% FBS. After 24 h, cells were infected with HSV-1 HF (range: 4×10^3^ - 4 PFU/mL) or KOS (2.5 PFUs/mL) and cultured with complete DMEM supplemented with 2% FBS and ACV 100 μM. After 5 days ACV is removed, and cells were maintained in complete DMEM with 0.5% FBS to monitor spontaneous reactivations. At 7 (HSV-1 HF) or 25 (HSV-1 KOS) days post infection (DPI), reactivation was induced using a thermal stress (1 h 30 min at 43°C) and cells were observed until reactivation occurred. 24 h after reactivation, cells were stained with Hoechst 33258 (Merck) and fixed with PFA/PBS 2%, or collected for total RNA extraction. Each condition was tested in quadruplicate. The fluorescence was measured by calculating mean gray values (average gray values within selected areas) using ImageJ software (NIH). Integrated density (area × mean gray value) was used to compare fluorescence intensities.

### Infection of DRGOs

For lytic infections, cell-free virus was adsorbed onto DRGOs seeded on Matrigel^®^ coated slides. After 2 h the cells were washed and cultured for 24 h, and live images were acquired. For latent infection, DRGOs were preincubated with ACV 100 μM in complete medium. After 24 h, DRGOs were infected with HSV-1 HF (range: 4×10^2^ - 4 PFU/mL) or KOS (2.5 PFU/mL) and cultured with complete medium with ACV 100 μM. After 5 days ACV is removed, and DRGOs were monitored for spontaneous reactivations. At 18 (HSV-1 HF) or 25 (HSV-1 KOS) DPI, reactivation was induced using a thermal stress (1 h 30 min at 43°C) and cells were observed until reactivation occurred. 24 h after reactivation, cells were stained with Hoechst 33258 (Merck) and fixed with PFA/PBS 2%, or collected for total RNA extraction. Each condition was tested in quadruplicate and integrated density was used to compare fluorescence intensities.

### Infection of microfluidic chip

Both NHEK and DRGOs were preincubated with ACV 100 μM for two days. Cells were infected with HSV-1 KOS (2.5 PFU/mL) and after 30 minutes of adsorption cells were washed and cultured with appropriate medium. Two experimental settings were performed, using 6 microfluidic chips for each condition (corresponding to 12 DRGOs) plus controls. For <u>experiment #1</u>, only DRGOs were infected and ACV 100 μM was added immediately after virus adsorption. For <u>experiment #2</u>, only NHEK were infected, and ACV 100 μM was added 6 hours after adsorption. After 5 days ACV is removed, and reactivation was induced using a thermal stress (1 h 30 min at 43°C). Two microfluidic chips (one for experiment) were not subjected to stimulus as control of spontaneous reactivation. Live images were acquired when reactivation occurred, and 24 hours later cells were collected for total RNA extraction. Integrated density was used to compare fluorescence intensities.

### ACV susceptibility

100 PFU/mL of virus were applied to Vero E6 monolayers. After 1h, virus mixture was replaced with complete DMEM with 2% FBS, 1% Agarose (BD) and serially diluted preparations of ACV (range: 0.75 – 400 μM). Plates were incubated for 46 h. Cells were then fixed and stained, and lysis plaques counted.

### Cell viability analysis

DRGOs were cocultured with HaCaT or NHEK on Matrigel^®^ coated slides, using different culture media: complete DRGOs media and complete DMEM with 10% FBS (50% v/v), complete DMEM with 10% FBS and Neurotrophins, complete DRGOs media and keratinocytes growth medium (50% v/v), keratinocytes growth medium and Neurotrophins. After 4 days, cytotoxicity was measured using *in vitro* toxicology assay kit, XTT based (TOX2; Merck)(Roehm et al. 1991) according to the manufacturer’s protocol. Incubation medium was collected after 4 h and read spectrophotometrically at a wavelength of 450 nm.

### Passive virus diffusion evaluation

Supernatants from chip chambers collected at different time points were applied to Vero E6 seeded on Matrigel^®^ coated 96 well Microplates μClear (Greiner Bio-One). Plates were centrifuged at 2,000 g for 15 minutes and incubated for 2 h. Then, supernatants were replaced with complete DMEM with 2% FBS and plates were incubated for 72 h. Live images were acquired after 48 h. Hoechst 33258 was used for nuclear staining before fixing cells with PFA/PBS 2%, 72 h post infection (PI).

### Immunofluorescence of DRGOs-NHEK connections

For immunocytochemical analysis, microfluidic chip were filled for 40 min at room temperature in 4% paraformaldehyde in PBS, washed with PBS at room temperature before permeabilized for 30 min in PBS containing 0.1% Triton X-100 and 10% normal goat serum and incubated overnight at 4 °C in PBS containing 10% normal goat serum and primary antibodies: anti-Cytokeratin 14 (K14) antibody (SP53; Merck), anti-Neurofilament heavy polypeptide (NF200) antibody (ab4680; Abcam), anti-Synapsyn 1 antibody (106001; Synaptic Systems), anti-VGluT1 antibody (ab77822; Abcam). Then cells were washed three times with PBS and incubated for 2 h at room temperature with secondary antibodies.

### Quantitative Two-step Reverse transcription-PCR (RT-qPCR)

Total RNA was extracted from pelleted cells with RNeasy Mini kit (74104; QIAgen) according to the manufacturer’s instructions. First-strand cDNA synthesis was performed on equivalent amounts of RNA from each sample, using an oligo(dT)15 primer and SuperScript IV Reverse Transcriptase (18091050; Thermo Fisher Scientific). Because the major species of latency-associated transcripts (LAT) is not polyadenylated (Farrell et al. 1991), reverse transcription of LAT was achieved by using a LAT-specific primer, ICP0-3’ (Tab.1, (Halford et al. 1996)). PCR was performed on equivalent amounts of cDNA; each reaction mixture contained 2 U Platinum Taq DNA polymerase (10966034; Thermo Fisher Scientific), 0.2 μM each PCR primer, 0.2 mM each deoxyribonucleotide triphosphate, PCR buffer and 1.5 mM MgCl2, and each reaction was performed in a MultiGene OptiMax (Labnet International) thermal cycler with 40 cycles of 94°C (30 s), 62°C (BRN3A, NAV1.7, ACTB) or 49°C (LAT) or 51°C (gG) or 55°C (GAPDH) or 59°C (gB) or 61°C (gD) (30 s), 72°C (30 s). Oligonucleotide primers were obtained from Eurofins Genomics and are listed in Table 1. Densitometry of GelRed-stained agarose gels was performed with Image-Lab software 6.0.1 (Bio-Rad). For each gene, the threshold at which product was detected was compared with the endogenous (GAPDH) control. Levels of viral mRNAs were normalized to GAPDH RNA levels to correct for recovery, because GAPDH levels at 6 hpi have been shown to be similar to levels in mock-infected cells (Schang et al. 1998).

**Table 1.**
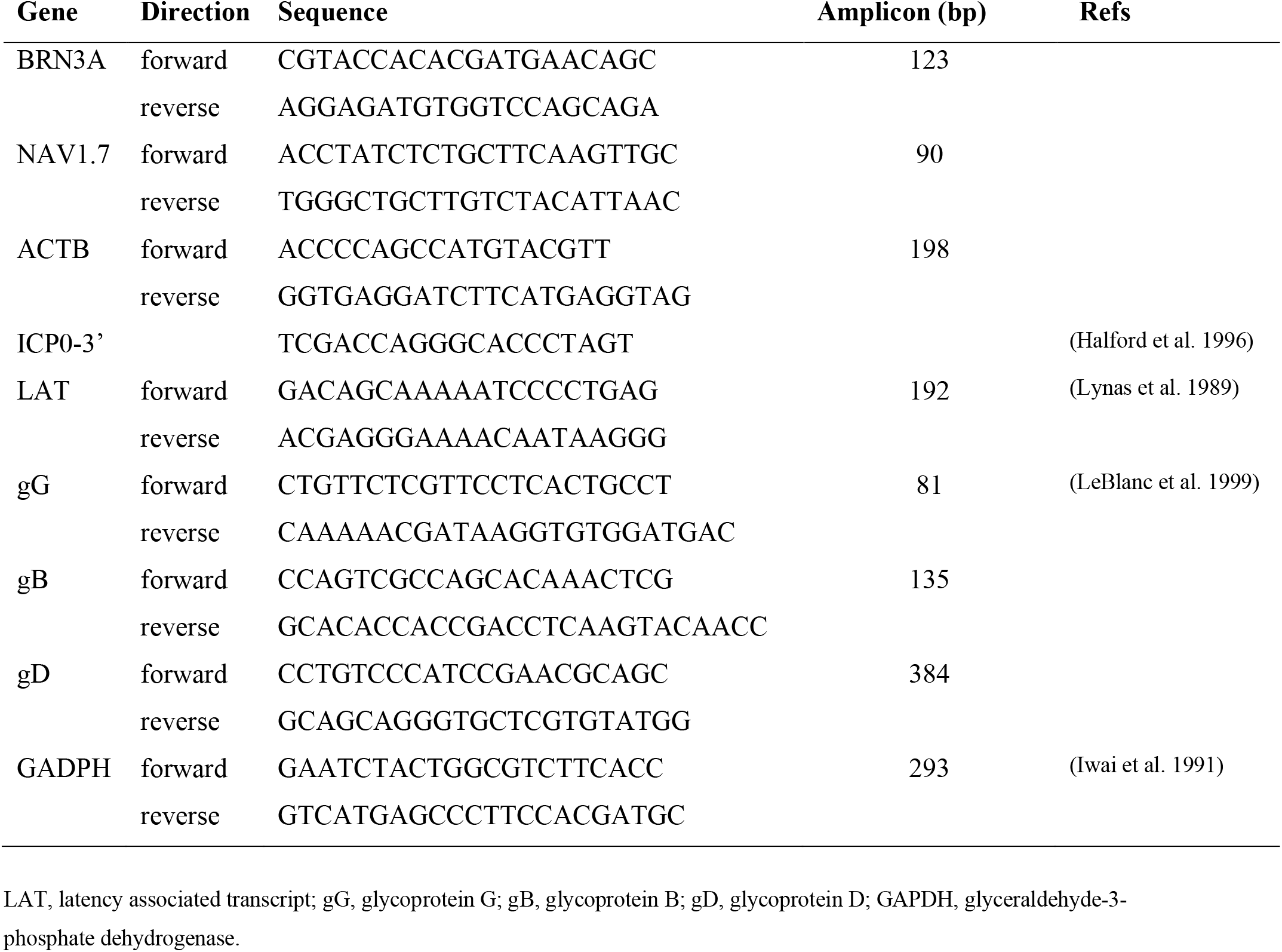
PCR primers and size of the amplicons.

### Statistical analysis

Two-way ANOVA and Sidak’s multiple comparisons test were performed for the evaluation of gene expression differences in mono-colure results and to compare fluorescence intensities in all experiments, while Tukey’s multiple comparisons test was used for the analysis of data from experiment #1 and #2.

## Data availability

Data generated during the study are available in the NCBI GEO public repository with accession n. GSE133755 (https://www.ncbi.nlm.nih.gov/geo/query/acc.cgi?acc=GSE133755) and GSE148212 (https://www.ncbi.nlm.nih.gov/geo/query/acc.cgi?acc=GSE148212).

## SUPPLEMENTARY FIGURES

**Supplementary Figure 1.**
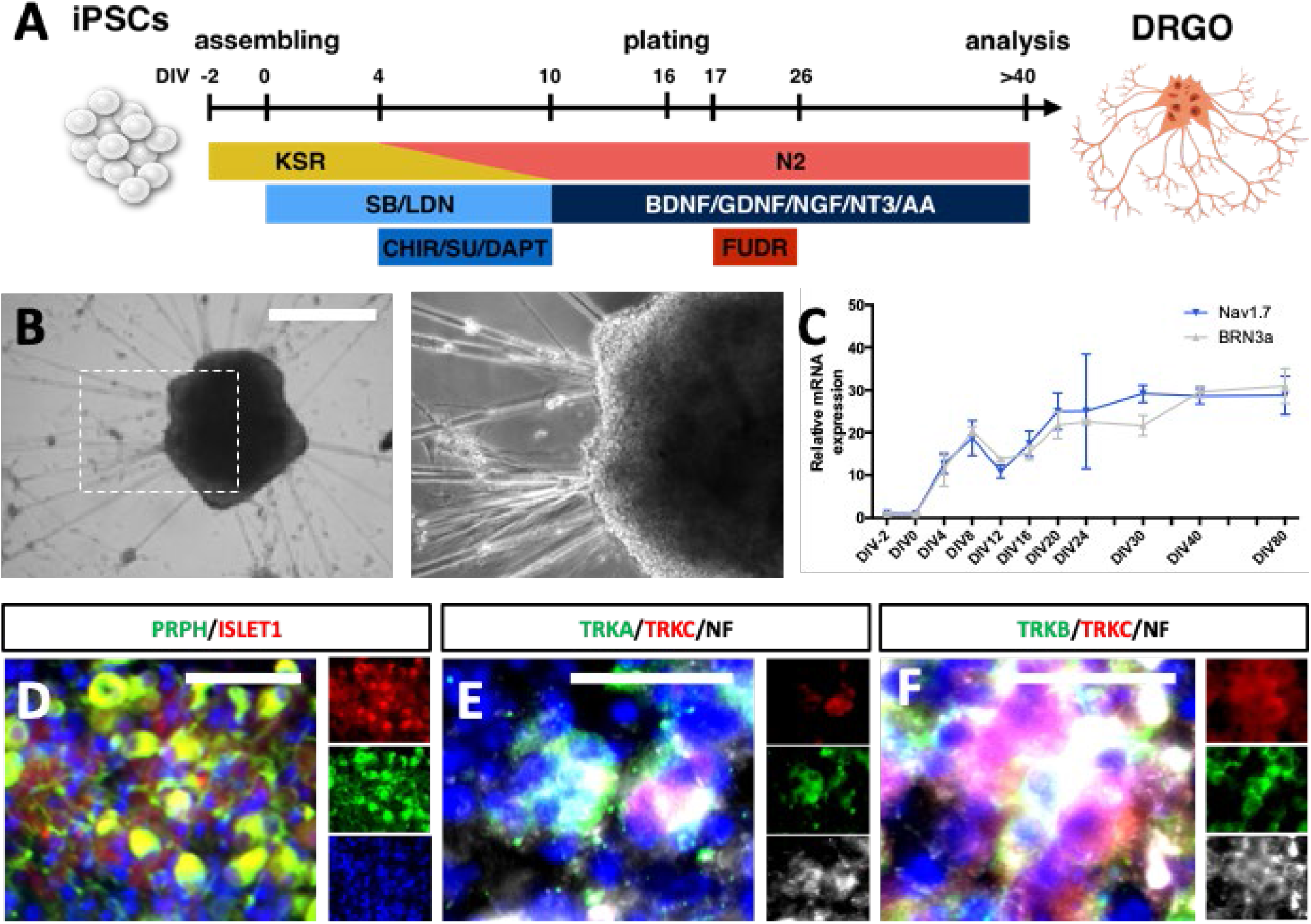
iPSC-derived organoid resembling primary dorsal root ganglia (DRGOs). **(A)** Illustration of the 3D culture system and sequential exposure to small molecules over time to obtain DRGOs. **(B)** Images of DRGO at DIV 40 with star-like web of neurites projections around the central mass and magnification. Scale bar, 500μm. (**C**) Quantitative analysis of *BRN3A* and *NAV1.7* transcript levels during DRGO differentiation. Expression levels are normalized to actin. Mean ± SD, n = 3 independent experiments, 8-12 organoids/line/experiment. **(D)** Immunocytochemistry in DRGOs at DIV 40 for proteins localized along neuronal projections peripherin, PRPH and the sensory neuron-specific transcription factors ISL1. Scale bar, 50μm. **(E)** Immunocytochemistry on DIV 40 DRGOs of Neurotrophic receptor tyrosine kinases (NTRK) 1/2/3 combination distinguishing the nociceptive (TRKA+/TRKC−). Scale bar, 10μm. **(F)** Immunocytochemistry to discriminate mechanoreceptive (TRKB+/TRKC+) and proprioceptive (TRKB-/TRKC+) neuronal subtypes. Scale bar, 10 μm.

**Supplementary Figure 2.**
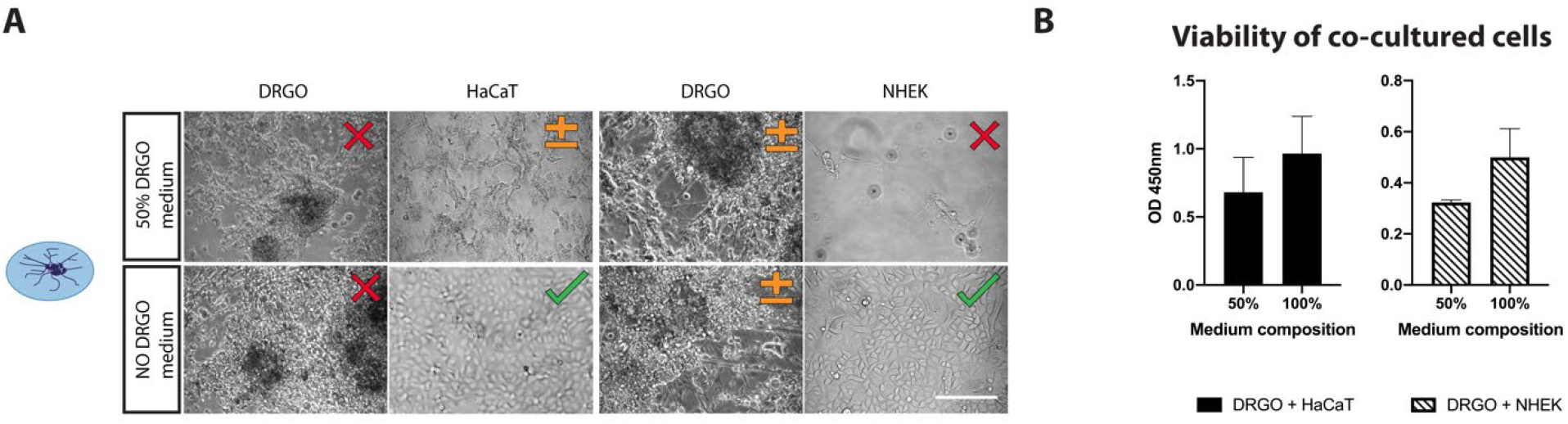
Unpattern co-culture of organoids and keratinocytes. (**A**) Bright-field microscopy images (20x magnification) of direct co-cultures of organoids (DRGOs) and both immortalized (HaCaT) and primary (NHEK) keratinocytes. Scale bar, 100 μm. **(B)** Cell proliferation assay to test DRGOs, HaCaT and NHEK viability. Mean + SD.

**Supplementary Figure 3.**
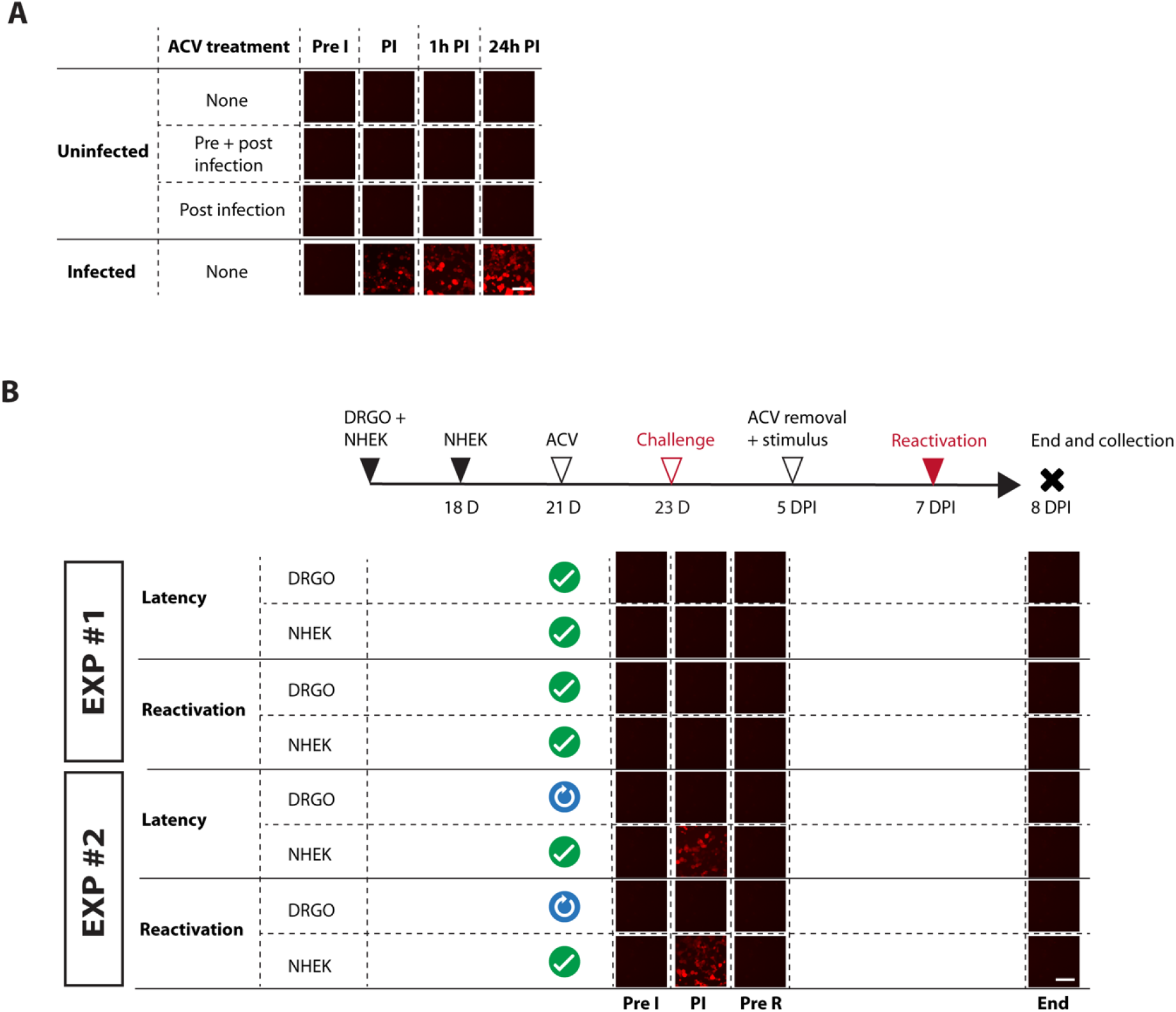
Microfluidics chip validation. **(A)** Chip supernatants collected at different time points tested on sensor cells for the presence of infective virus particles due to passive diffusion (lytic infection = red fluorescent signal). Scale bar, 500 μm. **(B)** Schematic representation of latency establishment and reactivation protocol used in chip EXP #1 and #2. Supernatants from different chambers and time points were tested on sensor cells as well (lytic infection = red fluorescent signal). Green tick indicates ACV addition to culture medium immediately after virus adsorption, blue arrow is for ACV delayed addition. (Pre I = before infection; PI = post-infection; Pre R = before reactivation). Scale bar, 500 μm, n = 6 independent experiments for each tested condition.

**Supplementary Figure 4.**
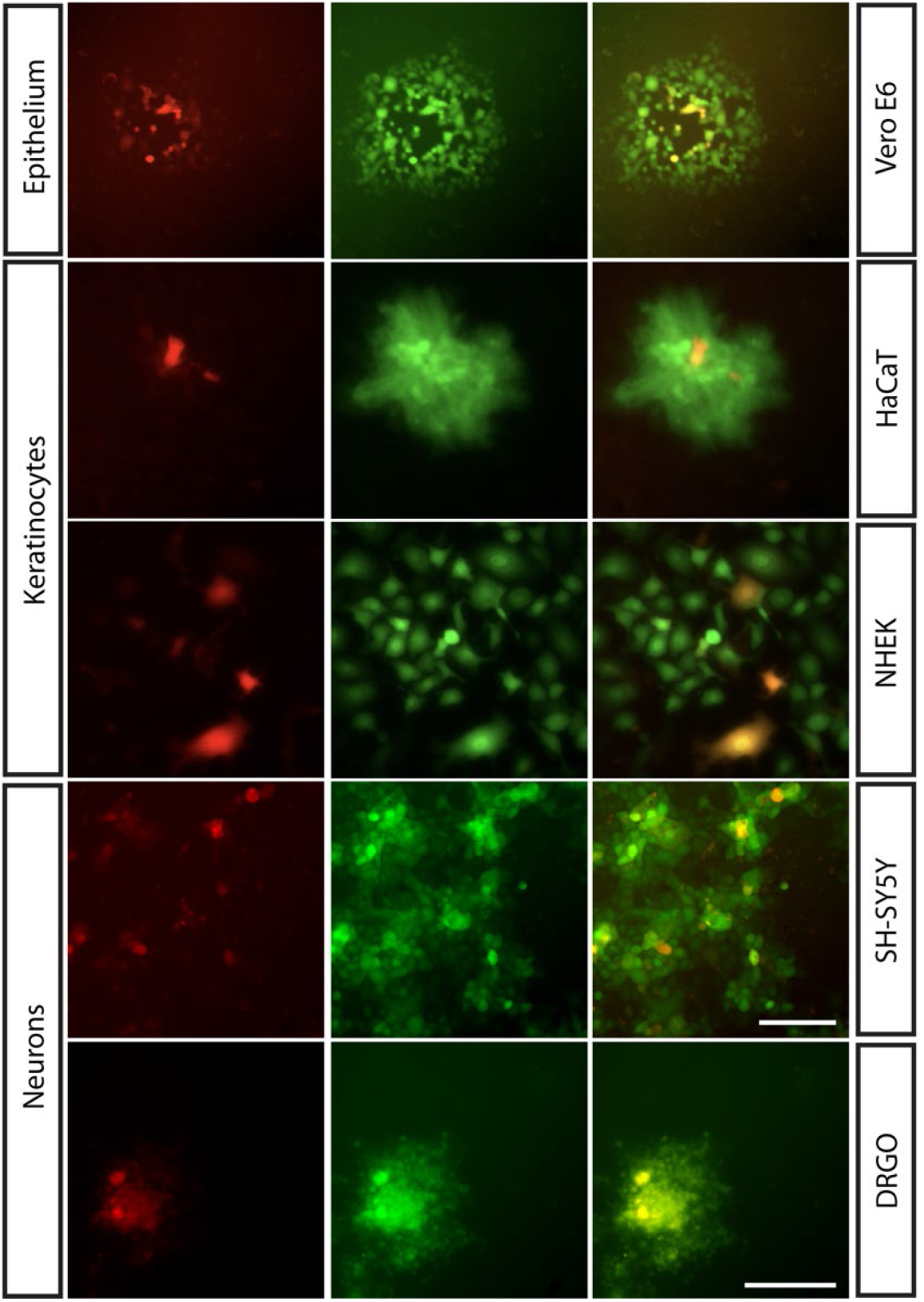
Lytic infection of different cell types using engineered HSV-1. Both immortalized (HaCaT) and primary keratinocytes (NHEK), neuroblastoma-derived cells (SH-SY5Y) and human iPSC-derived dorsal root ganglia organoids (DRGOs) were tested for their susceptibility to recombinant HSV-1 strain KOS infection. Epithelial cells (Vero E6) were used as a positive control of infection. EGFP (green, first column) and RFP (red, second column) signals indicate active gene expression driven by the viral promoters ICP0 and glycoprotein C, respectively. The last column shows the merge of the two fluorescent signals. Scale bar Vero E6/HaCaT/NHEK/SH-SY5Y 200 μm, DRGO 300 μm.

**Supplementary Figure 5.**
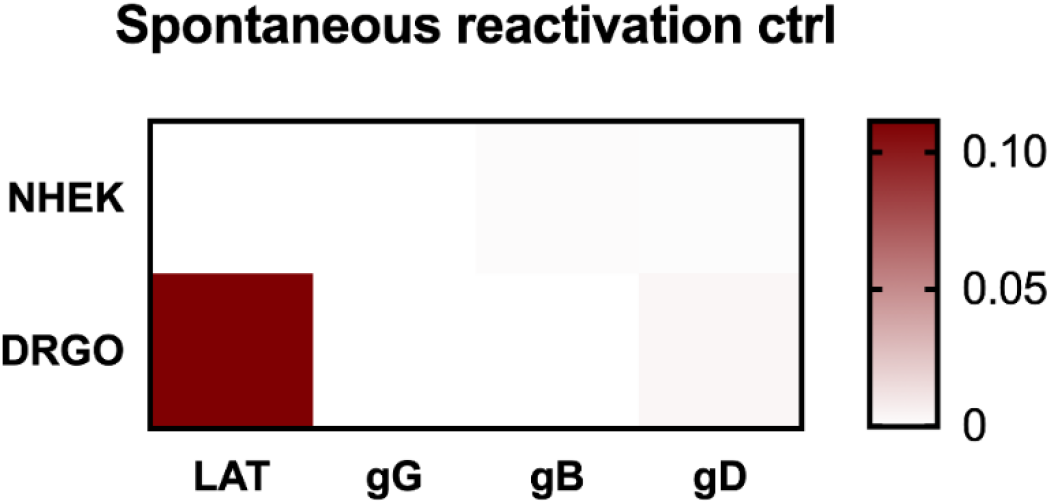
Control of spontaneous reactivation. Heatmap showing virus gene expression analysis of NHEK and DRGOs during latency without ACV addition to culture medium (dark red= high expression; white = no expression).

